# Swi/Snf Chromatin Remodeling Regulates Transcriptional Interference and Gene Repression

**DOI:** 10.1101/2023.04.27.538572

**Authors:** Kaitlin Morse, Sarah Swerdlow, Elçin Ünal

**Affiliations:** Department of Molecular and Cell Biology, Barker Hall, University of California, Berkeley, CA, USA, 94720

**Keywords:** gene regulation, transcription, transcriptional interference, Swi/Snf, chromatin, chromatin remodeling, LUTI, unfolded protein response

## Abstract

Alternative transcription start sites can affect transcript isoform diversity and translation levels. In a recently described form of gene regulation, coordinated transcriptional and translational interference results in transcript isoform-dependent changes in protein expression. Specifically, a long undecoded transcript isoform (LUTI) is transcribed from a gene-distal promoter, interfering with expression of the gene-proximal promoter. While transcriptional and chromatin features associated with LUTI expression have been described, the mechanism underlying LUTI-based transcriptional interference is not well understood. Using an unbiased genetic approach followed by integrated genomic analysis, we uncovered that the Swi/Snf chromatin remodeling complex is required for co-transcriptional nucleosome remodeling that leads to LUTI-based repression. We identified genes with tandem promoters that rely on Swi/Snf function for transcriptional interference during protein folding stress, including LUTI-regulated genes. To our knowledge, this study is the first to observe Swi/Snf’s direct involvement in gene repression via a *cis* transcriptional interference mechanism.

## INTRODUCTION

Gene regulation underlies proper development, homeostasis, and cellular stress response. Gene expression is initiated by transcription factors (TFs) and co-activators, which associate with promoters and enhancers to determine the timing, location, and amount of mRNA transcribed by RNA polymerase II (Pol II). According to classical models of gene regulation, transcript levels directly correlate with protein synthesis. However, the high prevalence of non-coding transcription and variation in translation efficiency among transcript isoforms has unveiled deeper complexity in the mRNA-to-protein relationship (Brar et al., 2012; Chia et al., 2021; Hangauer et al., 2013; Jorgensen et al., 2020; Pelechano et al., 2013; Wang et al., 2016).

Recently, an unconventional form of gene repression was discovered in yeast whereby mRNA and protein levels are inversely correlated for a given locus. For genes regulated in this manner, transcription initiation from a gene-distal promoter drives expression of a 5′ extended mRNA that contains the entire coding sequence (CDS) of the downstream gene (J. Chen et al., 2017; Cheng et al., 2018). However, upstream open reading frames (uORFs) in the 5′ leader of the extended mRNA restricts translation of the CDS. Based on these features, the distal mRNA isoform has been termed the long undecoded transcript isoform (LUTI). In addition to its translational repression, LUTI transcription interferes in *cis* with transcription of the CDS-proximal promoter, which controls expression of the coding mRNA isoform (Chia et al., 2017; Tresenrider et al., 2021). This coordinated transcriptional and translational interference by the LUTI reduces the pools of the coding mRNA in the cell by upregulating a coding-deficient mRNA isoform, ultimately resulting in decreased protein synthesis for the affected gene. Thus, LUTIs help explain cases in which mRNA and protein levels are poorly correlated (Cheng et al., 2018).

A key component of LUTI-based regulation is *cis*-mediated silencing of a canonical gene promoter by transcription of the LUTI. While numerous examples exist in which *cis*-acting noncoding RNAs regulate the expression of neighboring genes (Bird et al., 2006; Garg et al., 2018; J. H. Kim et al., 2016; Laprade et al., 2004; Werven et al., 2012), LUTI-based regulation distinctly involves the expression of an interfering mRNA transcript carrying a full CDS that is translationally repressed. In this regard, LUTI-based repression is counterintuitive from classical views of gene regulation as it involves TF activation of poorly translated mRNA isoforms, which ultimately leads to gene repression. Consequently, a single TF can coordinately activate or repress protein synthesis for distinct sets of genes, depending on whether it binds to a canonical or a LUTI promoter, respectively.

LUTI-mediated gene repression is pervasive. In yeast, hundreds of genes have an associated LUTI that is expressed at a distinct stage of meiosis to temporally restrict protein synthesis (Cheng et al., 2018). Additionally, cellular stress stimulates activation of LUTIs by the unfolded protein response (UPR) TF Hac1 and during zinc starvation by the zinc-responsive TF Zap1 (Van Dalfsen et al., 2018; Taggart et al., 2017). Case studies have revealed that diverse gene targets are subject to LUTI-based repression, including the genes encoding the kinetochore subunit Ndc80 (J. Chen et al., 2017; Chia et al., 2017), the superoxide dismutase enzyme Sod1 (Vander Wende et al., 2023), the transcription factor Swi4 (Su et al., 2023), and the purine hydrolase enzyme Hnt1 (Van Dalfsen et al., 2018; Tatip et al., 2020). Importantly, the observation that the human proto-oncogene *MDM2* is subject to LUTI-based interference has revealed this form of gene repression is broadly conserved (Hollerer et al., 2019).

Despite robust identification and characterization of LUTI-regulated genes, the mechanism underlying LUTI-based transcriptional interference is not well understood. In fact, transcript isoform profiling revealed that the degree of proximal promoter repression during LUTI expression is variable, indicating LUTI-mediated transcriptional repression is differentially regulated among affected loci (Tresenrider et al., 2021). Several factors are correlated with LUTI-based transcriptional interference in yeast, including high LUTI expression, increased histone 3 lysine 36 trimethylation (H3K36me3) at the proximal promoter, and changes in nucleosome positioning around the proximal promoter (Tresenrider et al., 2021). However, detangling causality from correlation for these chromatin and transcriptional features requires more in-depth functional analysis. In this study, we used an unbiased genetic approach to identify new regulatory factors required for LUTI-based transcriptional interference in budding yeast. This strategy led us to uncover a direct, repressive function of the Swi/Snf chromatin remodeling complex in establishing transcriptional interference.

The Swi/Snf complex is a highly conserved, twelve-subunit ATP-dependent nucleosome remodeling complex that activates the expression of a diverse set of genes in various contexts (Dutta et al., 2014; Rando and Winston, 2012; Rawal et al., 2018; Sahu et al., 2021; Shivaswamy and Iyer, 2008). We show that in addition to its canonical function in gene activation, the Swi/Snf complex performs nucleosome remodeling downstream of the active transcription start site (TSS) for its target loci. When the Swi/Snf complex is recruited to distal promoter targets, this downstream remodeling activity interferes with CDS-proximal promoters, leading to gene repression for select LUTI-regulated genes. In addition to furthering our understanding of LUTI-based transcriptional interference, our results clarify a long-standing question in the chromatin remodeling field by providing conclusive evidence that the Swi/Snf complex can directly repress transcription *in vivo* through its nucleosome remodeling activity.

## RESULTS

### A genetic approach to identify mutants defective in LUTI-based gene repression

To identify regulatory factors required for LUTI-based gene repression, we undertook a genetic approach. To create a reporter for LUTI regulation, we fused the 5′ leader sequence of a well-characterized LUTI, *NDC80^LUTI^*(J. Chen et al., 2017; Chia et al., 2017), to the CDS for the histidine biosynthesis gene *HIS3.* To enable inducible expression of the *HIS3^LUTI^* reporter, we replaced the native *NDC80^LUTI^* promoter with a *lexO* promoter, which can be induced by a β-estradiol activatable heterologous transcription factor (LexA-ER-B112) (Ottoz et al., 2014) (Figure 1A). We plated cells on media lacking histidine supplemented with β-estradiol to induce *HIS3^LUTI^* and 3-amino-1,2,4-triazole (3-AT) to completely inhibit low levels of His3 activity (refer to Materials and Methods for further details). These conditions should prevent the growth of cells where LUTI-based repression is intact (Figures 1A and 1B). However, disruption of the LUTI-based repression arising from spontaneous mutations should yield viable colonies under the same conditions due to loss of transcriptional and/or translational interference of *HIS3*. In fact, after three days of growth under selection, we observed viable colonies on the selective plates, which we termed “LUTI escape mutants.”

**Figure 1.**
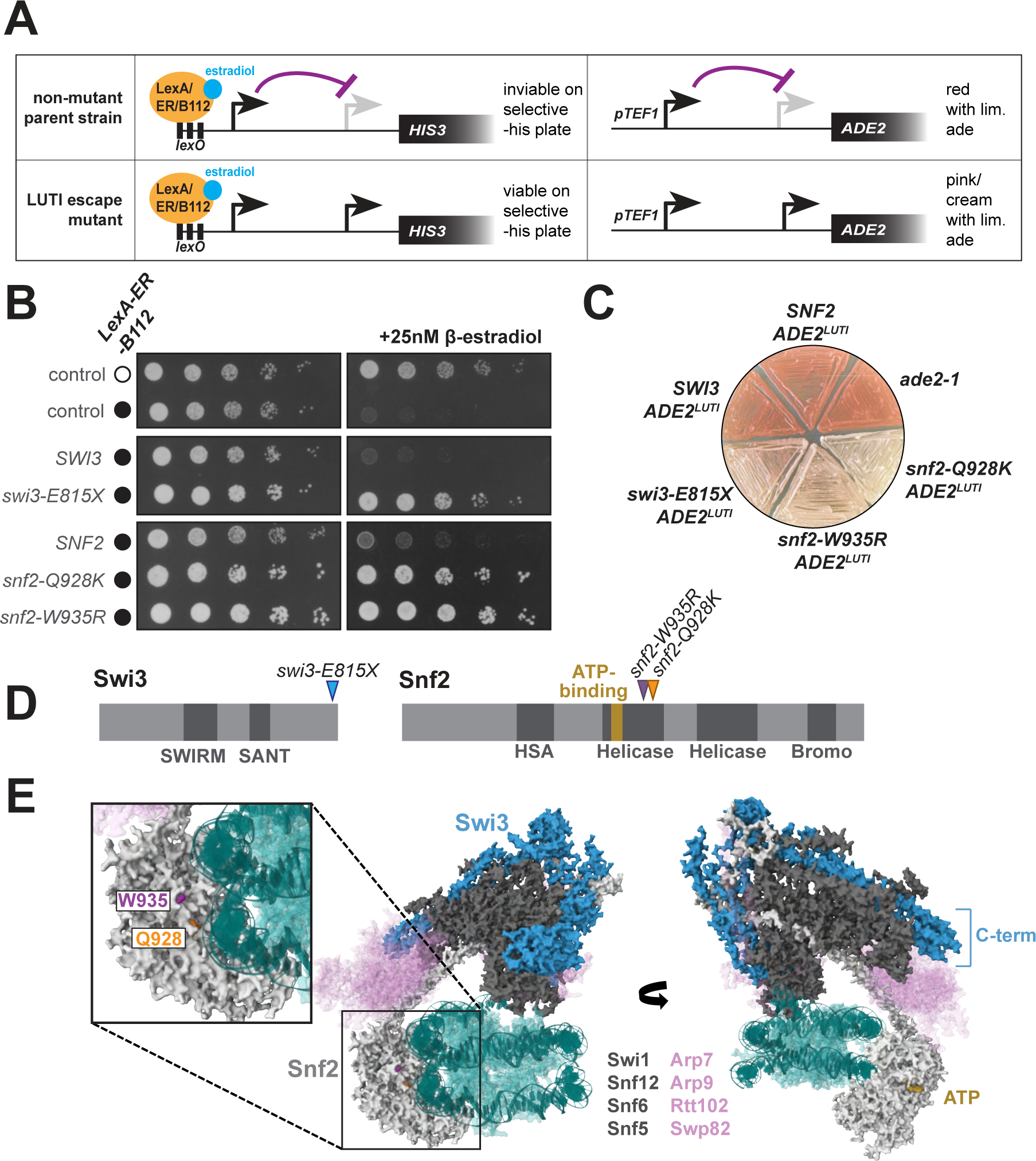
Mutations in the Swi/Snf complex disrupt LUTI-based transcriptional interference. **A.** Schematic of *HIS3^LUTI^* and *ADE2^LUTI^* reporters used to select for LUTI escape mutants. In control cells harboring the reporters (UB22912), expression of *HIS3^LUTI^* or *ADE2^LUTI^* silences expression of the protein-coding, proximal promoter-derived transcript, thereby rendering cells auxotrophic for histidine and adenine. In spontaneous LUTI escape mutants, cells that fail to silence *HIS3* expression can be selected based on their ability to grow in media lacking histidine, whereas cells that fail to silence *ADE2* expression can be screened based on their ability to metabolize red-pigmented purine precursors. **B.** *HIS3^LUTI^* serial dilution and spotting growth assay. Cells were plated on synthetic complete media lacking histidine with 200 µM 3-AT (left) or 200 µM 3-AT and 25 nM β-estradiol (right) and grown for 72 h at 30°C before imaging. Strains (from top to bottom): UB29385, UB29188, UB29791, UB24301, UB28911, UB28919, UB28925. **C.** *ADE2^LUTI^* color assay. Cells were streaked onto a YPD plate lacking supplemental adenine and grown for 24 h at 30°C before imaging. Strains: *ade2-1* (UB7), *SNF2 ADE2^LUTI^* (UB30034), *SWI3 ADE2^LUTI^* (UB30190), *swi3-E815X ADE2^LUTI^* (UB23545), *snf2-W935R ADE2^LUTI^* (UB28923), and *snf2-Q928K ADE2^LUTI^* (UB30185). **D.** Schematic of primary protein structure for Swi3 (left) and Snf2 (right), with LUTI escape mutations *swi3-E815X, snf2-W935R*, and *snf2-Q928K* mapped onto the structure (arrows). **E.** Cryo-EM structure of the yeast Swi/Snf complex bound to a nucleosome (teal), published by (Han et al., 2020) and rendered in MolStar (Sehnal et al., 2021). The Swi3 dimer is portrayed in blue, Snf2 in light gray, and other subunits with gene mutations recovered in the LUTI escape screen (Swi1, Snf12, Snf6, and Snf5) in dark gray. Other subunits (Arp7, Arp9, Rtt102, and Swp82) are portrayed in light pink. Snf11 was not resolved on this structure. The Snf2 residues W935 (purple) and Q928 (orange), which are affected by the *snf2-W935R* and *snf2-Q928K* mutations, are highlighted. The C-terminus of Swi3, which is affected by the *swi3-E815X* mutation, is highlighted however the specific region truncated by the *swi3-E815X* mutation is not resolved on this structure.

Mutants that we wished to filter out from our analysis were those that failed to express the LUTI in the first place, such as mutations that disrupt transcriptional activation that is dependent on the *lexO*/LexA-ER-B112 inducible system. Accordingly, we performed secondary screening on each mutant using an independent reporter. For this, we used a constitutive, highly expressed *TEF1* promoter to drive LUTI expression and replaced the *HIS3* CDS with the CDS for the adenine biosynthesis gene *ADE2* (Figure 1A). In this case, cells that properly silenced *ADE2* expression turned red on media with limited adenine due to their accumulation of red-pigmented purine precursors, whereas LUTI escape mutants appeared pink or cream-colored due to their failure to repress *ADE2* (Figure 1C).

To identify the causative mutations behind the LUTI escape phenotypes, we then sequenced the genomes of cells exhibiting both *HIS3^LUTI^*and *ADE2^LUTI^* escape phenotypes. We identified and validated eleven mutations conferring LUTI suppressor phenotypes (see Materials and Methods for details). Strikingly, all identified LUTI escape mutations fell within genes encoding subunits of the Swi/Snf chromatin remodeling complex (Table S1, Figure 1E). Our selection-based strategy uncovered mutations in six of the eight subunits that are specific to the Swi/Snf complex and are not members of other chromatin remodeling complexes (Olave et al., 2002; Peil et al., 2022; Turegun et al., 2018), with five of the identified mutations falling within the gene encoding the catalytic subunit, *SNF2* (Table S1).

We did not identify mutations in *SWP82* or *SNF11*, which also encode subunits specific to the Swi/Snf complex. We investigated whether deletion of these genes would disrupt LUTI*-*based repression and found that the *swp82Δ* mutation did not confer a LUTI escape phenotype for the *HIS3^LUTI^*or *ADE2^LUTI^* reporters (Figures S1A and S1B). This may be due to a limited role for *SWP82* in Swi/Snf function as judged by the lack of a growth defect in the null mutant (Figure S1C). Furthermore, *swp82Δ* was previously shown to confer milder gene expression defects compared to other null mutants of Swi/Snf (Dutta et al., 2017). Unlike *swp82Δ*, *snf11Δ* cells did exhibit subtle LUTI escape phenotypes (Figures S1A and S1B). Snf11 is only 169 amino acids long and may have eluded our selection approach, which relied on spontaneous mutations, due to its small size.

It was surprising that we only recovered mutations in the subunits of Swi/Snf and no other chromatin remodelers. In fact, Chd1 and Isw-family chromatin remodelers have known functions in co-transcriptional nucleosome remodeling (Hennig et al., 2012; Rando and Winston, 2012; Smolle et al., 2012). To assess their involvement in LUTI-based repression, we deleted catalytic subunits from each complex. We found that deletion of *ISW1, ISW2,* or *CHD1*, alone or in combination, did not disrupt *HIS3^LUTI^* or *ADE2^LUTI^*-based repression (Figures S1D and S1E). This finding suggests that the Swi/Snf complex plays a specific role in LUTI-based gene repression for the *HIS3^LUTI^* and *ADE2^LUTI^* reporter genes.

### LUTI escape mutations confer partial loss of Swi/Snf function

All LUTI escape mutants conferred recessive phenotypes except for *snf2-Q928K* (Table S1), which displayed a partial dominant phenotype (Table S1 and Figure 2A). We chose to use this mutant along with *snf2-W935R* and *swi3-E815X*, a nonsense mutant affecting the structural subunit Swi3, to further investigate the role of Swi/Snf chromatin remodeling in transcriptional interference (Figures 1D and 1E). These three mutants were selected for further characterization based on their minimal growth defects compared to *snf2Δ* or *swi3Δ* (Figure S2A) and strong LUTI escape phenotypes (Figures 1B and 1C). For each gene, we constructed strains lacking the endogenous allele and harboring a transgenic rescue construct at the *LEU2* locus containing either the wild-type allele of the gene as a control or the LUTI escape allele, under the native gene promoter.

**Figure 2.**
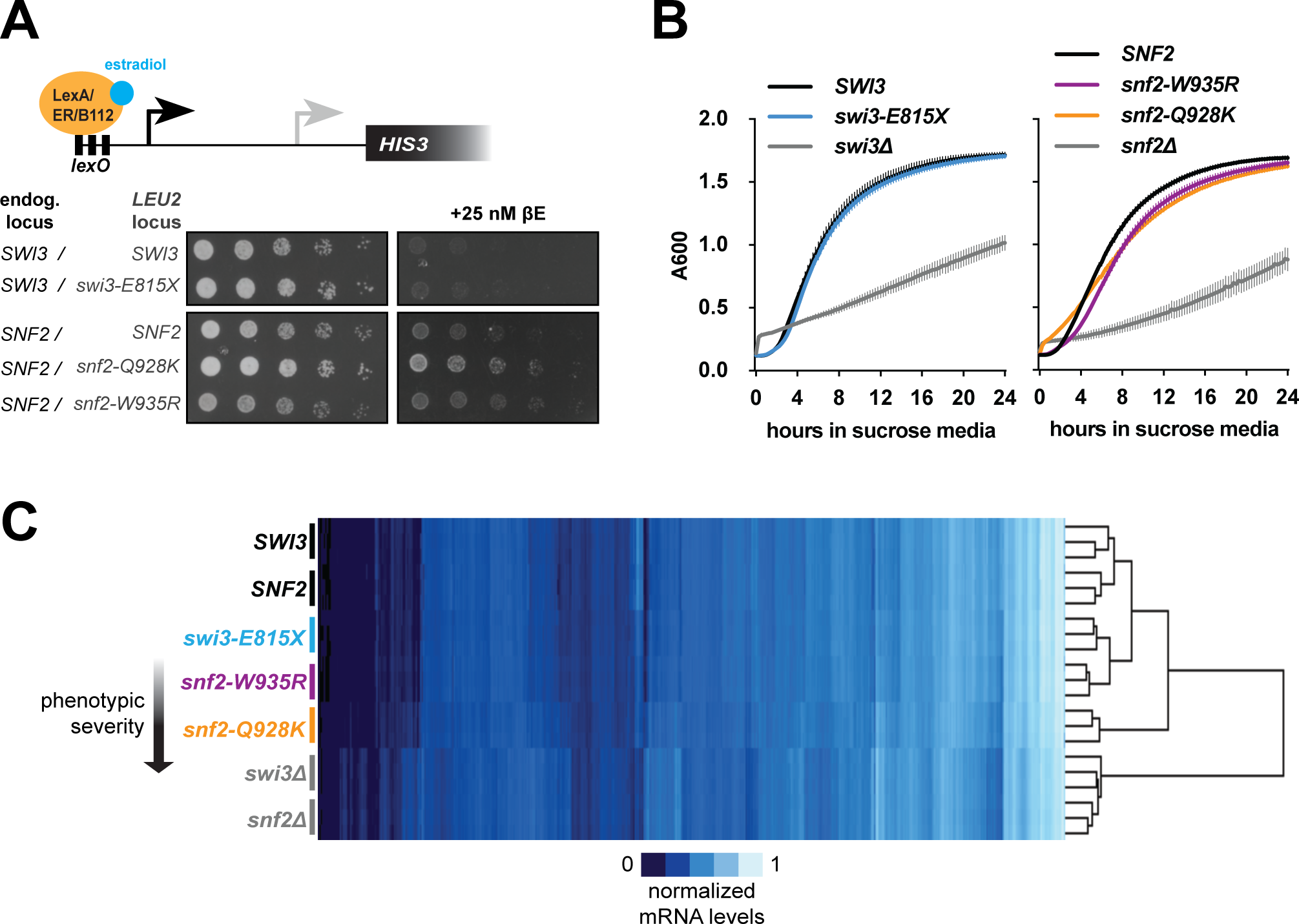
LUTI escape mutations confer partial loss of Swi/Snf function. **A.** *HIS3^LUTI^* serial dilution and spotting growth assay in haploid cells harboring transgenic alleles at the LEU2 locus for *SWI3* (UB29792), *swi3-E815X* (UB29694), *SNF2* (UB28907), *snf2-Q928K* (UB29170), or *snf2-W935R* (UB29166). **B.** Growth curves for cells grown in rich media with 2% sucrose. Absorbance readings at 600 nm collected at 15-minute intervals for 24 h is plotted for *SWI3* (UB19205, left, black), *swi3-E815X* (UB19209, left, blue), *swi3Δ* (UB27896, left, gray), *SNF2* (UB28914, right, black), *snf2-Q935R* (UB28922, right, purple), *snf2-Q928K* (UB28915, right, orange), and *snf2Δ* (UB29781, right, gray). **C.** Heatmap of hierarchical clustering performed on genes (x-axis, Euclidian-distance similarity metric) and strains in biological triplicate (y-axis, centered correlation similarity metric) produced from mRNA sequencing TPM values. Strains (top to bottom) are the same as listed in (B). All three LUTI escape mutants cluster more closely with wild-type controls than the null mutants, however the *snf2-Q928K* mutant has a gene expression profile that is more divergent from wild type than the *swi3-E815X* or *snf2-W935R* mutants.

Because null mutations in *SNF2* or *SWI3* are extremely pleiotropic (Dutta et al., 2017), we wondered whether the LUTI escape mutants broadly share phenotypes with their respective null mutants or if they instead affect specific functions of the Swi/Snf complex. To address this question, we first examined a well-characterized Swi/Snf loss-of-function phenotype: inability to ferment sucrose (Neigeborn and Carlson, 1984). As expected, *snf2Δ* and *swi3Δ* mutants grew poorly in sucrose media (Figure 2B). In contrast, the *swi3-E815X* mutant grew at a rate identical to wild-type cells, while the *snf2-W935R* and *snf2-Q928K* mutants only exhibited slight growth defects (Figure 2B).

To better understand how the LUTI escape mutants affect gene expression at a global scale, we next performed mRNA-seq on *swi3-E815X*, *snf2-W935R,* and *snf2-Q928K* mutants along with the respective wild-type control and null mutant. While the *snf2Δ* and *swi3Δ* mutants displayed widespread changes in gene expression compared to wild-type cells (Spearman’s rank correlation coefficient, ρ = 0.883 [*snf2Δ*], ρ = 0.892 [*swi3Δ*]; Figures 2C and S2D), the LUTI escape mutants affected only a limited number of genes (Figure 2C). Hierarchical clustering further revealed that *swi3-E815X* and *snf2-W935R* mutants were grouped with the wild-type controls and displayed gene expression profiles nearly matching that of wild-type cells (ρ = 0.98 [*swi3-*E815X], ρ = 0.97 [*snf2-W935R*]; Figure S2D). In contrast, the *snf2-Q928K* mutant displayed an intermediate gene expression profile, with some genes matching the null phenotype and some matching wild-type gene expression (ρ = 0.947 [*snf2-Q928K* vs. wild type], ρ *=* 0.95 [*snf2-Q928K* vs. *snf2Δ*]; Figures 2C and S2D). These findings confirm that the *snf2-Q928K* mutant displays more severe and pleiotropic loss-of-function phenotypes compared to *snf2-W935R* and *swi3-E815X*.

The Swi/Snf complex has been previously implicated in transcriptional interference of the serine biosynthesis gene *SER3*. In serine-rich conditions, Swi/Snf activates an upstream intergenic non-coding RNA called *SRG1*, which reads through the *SER3* promoter resulting in increased nucleosome occupancy and repression of *SER3* (Hainer et al., 2011; Martens et al., 2005; Martens and Winston, 2002). In YPD, a serine-rich media, both *snf2Δ* and *swi3Δ* cells exhibited lower levels of *SRG1* transcript compared to wild-type cells (Figure S2B), leading to significant upregulation of *SER3* (p= 0.0058 [*snf2Δ*], p= 0.0018 [*swi3Δ*], Figure S2C). The *snf2-Q928K* mutant downregulated *SRG1* transcript to the same degree as *snf2Δ* cells (Figure S2B) and upregulated *SER3* ∼40-fold relative to wild type (Figure S2C), whereas the *swi3-E815X* and *snf2-W935R* mutants regulated this locus normally. Thus *snf2-W935R* and *swi3-E815X* mutants appear to display defects related to transcriptional interference at specific loci. Altogether, we conclude that each LUTI escape mutation disrupts Swi/Snf function, but to varying degrees, with *snf2-Q928K* exhibiting more severe transcriptional defects than *snf2-W935R*, and *snf2-W935R* exhibiting slightly more defects than *swi3-E815X* (Figure 2C).

### Swi/Snf regulates DTT-induced alternative transcript isoform expression

Although Swi/Snf has been shown regulate transcriptional interference of *SER3* indirectly through activation of the interfering transcript *SRG1,* direct interference through Swi/Snf chromatin remodeling at a silenced promoter has not previously been observed. To explore this activity on a genome-wide scale, we turned to a cellular context during which transcript isoform toggling is widespread: the unfolded protein response (UPR) (Van Dalfsen et al. 2018). In order to quantify differences in transcript isoform expression for genes with alternative transcription start sites (TSSs), we performed transcript leader sequencing (TL-seq) (Arribere and Gilbert, 2013) in wild type and LUTI escape mutants that were either untreated or treated with dithiothreitol (DTT) to induce the UPR. To restrict our analysis to loci in which the distal TSS (TSS^DIST^) drives readthrough transcription across the CDS-proximal TSS (TSS^PROX^), rather than cases of intergenic transcription that terminates upstream of the proximal promoter, we also performed Nanopore direct mRNA-sequencing (direct mRNA-seq) in wild-type cells to visualize full-length mRNA isoforms. Finally, we excluded indirect gene targets that do not exhibit Snf2 binding with DTT treatment by performing Snf2 chromatin immunoprecipitation followed by whole genome sequencing (ChIP-seq) (Figure 3A).

**Figure 3.**
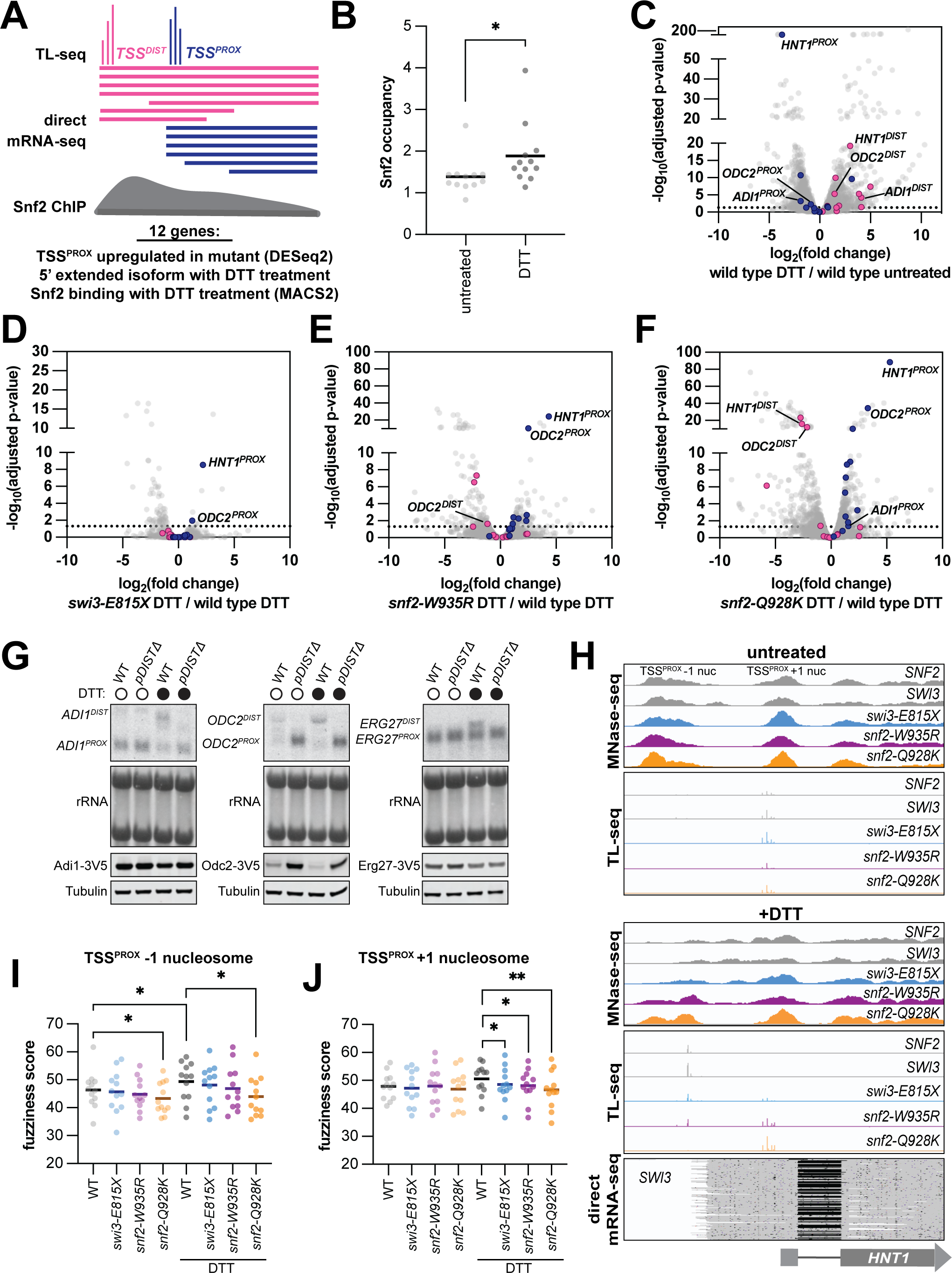
Swi/Snf regulates DTT-induced alternative transcript isoform expression. **A.** Schematic of the strategy to identify Swi/Snf-regulated genes that exhibit transcript toggling upon UPR induction by treatment with 5 mM DTT for 1 h. Twelve genes were identified that (1) exhibited significant upregulation in one or more Swi/Snf mutant of the TSS^PROX^ (DESeq2, p < 0.05), were (2) subject to transcriptional readthrough from an upstream distal promoter and (3) have a Snf2 ChIP peak that was called by MACS2. **B.** Snf2 ChIP-seq signals plotted for the 12 genes identified by the strategy outlined in (A). The average Snf2 binding levels in wild-type cells (UB30387 and UB30070, n = 2 for each wild-type *SNF2* and *SWI3* control strain, see Materials and Methods for details) is 1.4-fold higher on average when cells are treated with DTT compared to the unstressed condition (paired t test, two-tailed, p = 0.0423). **C.** Volcano plot produced from the output of differential gene expression analysis (DESeq2) on TL-seq data for wild-type cells (UB19205 and UB28914, see Material and Methods for details) that were untreated or treated with DTT (n = 4). Distal TSSs (pink) and proximal TSSs (dark blue) for genes that are Swi/Snf regulated and exhibit promoter toggling with stress induction are highlighted. **D-F.** Same as (C), but comparing **(D)** DTT-treated *swi3-E815X* (UB19209) cells to DTT-treated *SWI3* (UB19205) cells (n = 2), **(E)** DTT-treated *snf2-W935R* (UB28922) cells to DTT-treated *SNF2* (UB28914) cells (n = 2), and **(F)** DTT-treated *snf2-Q928K* (UB28915) cells to DTT-treated *SNF2* (UB28914) cells (n = 2). **G.** RNA blot (top) and immunoblot (bottom). Both blots are specific for the V5 sequence that is C-terminally fused to *ADI1* (left), *ODC2* (middle), or *ERG27* alleles integrated at the *TRP1* locus. Transgenes either harbored the TSS^DIST^ and its promoter (WT) or lacked this sequence (*pDISTΔ*). Untreated control cells or cells treated with 5 mM DTT were collected one h post-induction. rRNA is detected on the RNA blot by methylene blue staining. For the immunoblot, alpha-tubulin is used as a loading control. One of two biological replicates is shown. Strains: *ADI1* (UB36511), *ADI1^pDISTΔ^* (UB36513), *ODC2* (UB36515), *ODC2^pDISTΔ^* (UB36521), *ERG27* (UB36594), and *ERG27^pDISTΔ^* (UB36596). **H.** Genome browser snapshots portraying MNase-seq, TL-seq, and Nanopore direct mRNA-seq for the *HNT1* locus for *SNF2* (UB30387), *SWI3* (UB30070), *swi3-E815X* (UB30071), *snf2-W935R* (UB30391), and *snf2-Q928K* (UB30389) cells that were untreated (top) or treated with 5 mM DTT (bottom). One of two biological replicates is portrayed, except for direct mRNA-sequencing for which only one replicate was performed. **I.** Nucleosome fuzziness scores output from DANPOS3 for the -1 nucleosome relative to the TSS^PROX^ in wild-type cells (n = 4, strains UB30387 and UB30070) and *swi3-E815X* (UB30071), *snf2-W935R* (UB30391), and *snf2-Q928K* (UB30389) mutants (n = 2) that were untreated or treated with 5 mM DTT. A paired t test was performed on wild type fuzziness scores comparing untreated to DTT-treated cells (two-tailed, p = 0.0462) and each mutant-to-wild type comparison in untreated conditions (p = 0.5938 [*swi3-E815X*], p = 0.2487 [*snf2-W935R*], p = 0.0371 [*snf2-Q928K*]) or with DTT treatment (p = 0.1283 [*swi3-E815X*], p = 0.2043 [*snf2-W935R*], p = 0.0129 [*snf2-Q928K*]). **J.** Same as (I), but for the +1 nucleosome relative to the TSS^PROX^. A paired t test was performed on wild type fuzziness scores comparing untreated to DTT-treated cells (two-tailed, p = 0.0729) and each mutant-to-wild type comparison in untreated conditions (p = 0.4463 [*swi3-E815X*], p = 0.9578 [*snf2-W935R*], p = 0.4348 [*snf2-Q928K*]) or with DTT treatment (p = 0.0470 [*swi3-E815X*], p = 0.0397 [*snf2-W935R*], p = 0.0040 [*snf2-Q928K*]).

We uncovered 12 TSS^PROX^ loci that fit the following criteria upon DTT treatment: (1) The TSS^PROX^ was significantly upregulated in one or more of the LUTI escape mutants (DESeq2, adjusted p-value > 0.05); (2) A TSS^DIST^-driven readthrough transcript was expressed; and (3) Snf2 was enriched at the corresponding locus. Upon DTT-dependent induction of TSS^DIST^ transcription, Snf2 occupancy levels increased at the 5′ regulatory region by an average of 1.4-fold compared to unstressed conditions (paired t test, two-tailed, p = 0.0423; Figure 3B). Excitingly, the TSS^PROX^ for the previously characterized LUTI-regulated gene *HNT1* (Van Dalfsen et al. 2018) was significantly upregulated in all three LUTI escape mutants (Figures 3D-F). Furthermore, analysis of a previously published ribosome profiling dataset (Van Dalfsen et al. 2018) revealed that *ADI1* and *ODC2* also exhibited uORF translation in the 5′ leader sequence of their distal mRNA isoform (Figure S3B). Deletion of the distal promoter for *ADI1* and *ODC2* resulted in increased abundance of the TSS^PROX^-derived mRNA isoform and increased protein levels (Figures 3G and S3C), revealing these are also LUTI-regulated genes subject to both transcriptional and translational interference upon DTT treatment.

For nine of the 12 genes, the distal mRNA isoform did not fit the criteria of a LUTI because it did not display evidence of lower translation efficiency compared to the proximal isoform (Table S2). However, these genes remain useful models to investigate Swi/Snf-dependent transcriptional interference, given that LUTI-based transcriptional interference and translational repression are not mechanistically coupled. For these non-LUTI cases, Swi/Snf may act to repress the TSS^PROX^ upon TSS^DIST^ activation, however there is no corresponding impact on protein levels. Indeed, when we deleted the distal promoter for *ERG27,* a gene that did not exhibit isoform-dependent protein level changes (Figure S3C), there was a subtle increase in *ERG27^PROX^* expression (Figure 3G). We conclude that the Swi/Snf complex represses transcription at select promoters that are subject to transcriptional readthrough upon DTT treatment, including LUTI-regulated promoters.

Among the LUTI escape mutants, the *snf2-Q928K* mutant displayed the most changes in genome-wide TSS expression levels relative to wild type (Figure 3F). This is consistent with mRNA-seq results which revealed this mutant was more pleiotropic compared to the *snf2-W935R* or *swi3-E815X* mutants. In *snf2-Q928K* cells, the TSS^DIST^ for five genes was downregulated (*HNT1, ODC2, PRY1, ERG27,* and *FLC1*), thus TSS^PROX^ upregulation for these genes likely results from reduced transcriptional readthrough in this mutant (Figure 3F, Table S2). The *snf2-W935R* mutant also exhibited downregulation of the TSS^DIST^ for four genes (*ODC2, PRY1, ERG27,* and *FLC1*), albeit to a lesser extent than in *snf2-Q928K* cells, consistent with the *snf2-W935R* mutation conferring more mild loss of Swi/Snf function compared to the *snf2-Q928K* mutation (Figure 3E). Finally, the *swi3-E815X* mutant had the fewest changes in TSS expression compared to wild type, both globally and among the 12 transcriptional interference targets, exhibiting significant upregulation of the *ODC2^PROX^*and *HNT1^PROX^* isoforms and no significant changes for TSS^DIST^ levels (Figure 3D).

Overall, TL-seq analyses reveal that the Swi/Snf complex regulates transcriptional interference in response to protein folding stress through two routes: activation of distal promoters and repression of the downstream proximal promoters. It seems the *snf2-Q928K* mutation reduces canonical transcription initiation activity by the Swi/Snf complex, consistent with our previous finding that *SRG1* expression is reduced in this mutant. However, the *snf2-W935R* mutation only slightly reduces initiation of the TSS^DIST^ for some loci, yet still disrupts repressive activity by Swi/Snf at the TSS^PROX^. The *swi3-E815X* mutation also reduces repressive activity by Swi/Snf at the TSS^PROX^ but to a lesser extent than the *snf2-W935R* mutant.

### Nucleosome remodeling is reduced at the TSS^PROX^ in Swi/Snf LUTI escape mutants

To investigate whether transcriptional interference at loci affected by LUTI escape mutants is mediated by changes in chromatin structure, we performed micrococcal nuclease digestion and whole genome sequencing (MNase-seq). When we analyzed the nucleosome profiles for the LUTI-regulated gene *HNT1* in wild-type cells, we observed a shift from stable nucleosome positioning for the -1 and +1 nucleosomes surrounding the *HNT1^PROX^*TSS to fuzzy positioning upon DTT treatment (Figure 3H), indicating that *HNT1^LUTI^*expression is associated with nucleosome remodeling downstream of the *HNT1^LUTI^*TSS. In contrast, the +1 nucleosome relative to the *HNT1^PROX^* TSS remained stably positioned in all three LUTI escape mutants (Figure 3H), suggesting LUTI-coupled nucleosome remodeling is impaired in these mutants. The -1 nucleosome relative to the *HNT1^PROX^* TSS, which spans the *HNT1^LUTI^* TSS in untreated cells, also displayed a stronger MNase-seq signal in the *snf2-Q928K* and *snf2-W935R* mutants with DTT treatment, although its position was shifted compared to the untreated condition (Figure 3H). Lack of remodeling for this nucleosome may preclude *HNT1^LUTI^* expression during UPR induction in *snf2-Q928K* cells, as remodeling of the chromatin upstream of the *HNT1^LUTI^* TSS is associated with high activation of *HNT1^LUTI^* in wild-type cells.

We next investigated whether nucleosome positioning was also stabilized in the LUTI escape mutants for the other 11 TSS^PROX^ loci. We compared nucleosome fuzziness scores, a quantitative measurement for nucleosome positioning in which higher fuzziness corresponds to more poorly positioned nucleosomes, across wild-type and mutant cells. Both the -1 and +1 nucleosomes surrounding each TSS^PROX^ became more fuzzy in wild-type cells upon DTT treatment compared to unstressed conditions, indicating remodeling of these nucleosomes is associated with distal promoter expression (paired t test, two-tailed, p = 0.0462 [-1 nuc], p = 0.0729 [+1 nuc]; Figures 3I and 3J). For half of the 12 genes investigated, as with *HNT1*, the TSS^PROX^ -1 nucleosome is also the +1 nucleosome for the TSS^DIST^. In these cases, remodeling of this nucleosome may be required to facilitate proper initiation of the distal isoform. Nucleosome fuzziness for the TSS^PROX^ -1 nucleosome was not significantly different in the *swi3-E815X* or *snf2-W935R* mutants compared to wild type (p = 0.1283 [*swi3-E815X*], p = 0.2043 [*snf2-W935R*]), but there was a significant decrease in fuzziness for the *snf2-Q928K* mutant (p = 0.0129; Figure 3I). Together, these results combined with our TL-seq results are consistent with a model in which *snf2-Q928K* cells are defective for transcription initiation, as opposed to *snf2-W935R* and *swi3-E815X* cells, which may be defective for nucleosome remodeling at positions downstream from the transcriptionally active TSS^DIST^.

In response to protein folding stress, all three LUTI escape mutants exhibited reduced nucleosome fuzziness for the TSS^PROX^ +1 nucleosome compared to wild-type cells (p = 0.0470 [*swi3-E815X*], p = 0.0397 [*snf2-W935R*], p = 0.0040 [*snf2-Q928K*]; Figure 3J). Excitingly, these results align with the previous finding that increased TSS^PROX^ +1 nucleosome fuzziness was found to be correlated with more potent LUTI-based transcriptional interference (Tresenrider et al., 2021). We also examined changes in nucleosome occupancy within the nucleosome depleted region (NDR) for the TSS^PROX^ and found that upon DTT treatment, nucleosome occupancy within the NDR increased on average by 1.9-fold compared to the unstressed condition (p = 0.0445; Figure S3D). Each LUTI escape mutant had lower nucleosome occupancy within the TSS^PROX^ NDR compared to wild-type cells upon DTT treatment (not significant, Figure S3D), indicating that increased expression of proximal mRNA isoforms in the mutants may result from increased promoter accessibility.

### Swi/Snf facilitates rapid and sustained repression of *HNT1^PROX^* upon *HNT1^LUTI^* induction

To further dissect the role of the Swi/Snf complex in transcriptional interference, we next examined the kinetics of *HNT1^PROX^* repression and chromatin changes when *HNT1^LUTI^* is induced. Wild-type cells induced *HNT1^LUTI^*within 5 minutes of DTT treatment and expression of *HNT1^LUTI^* further increased 3-fold by 30 minutes, at which time *HNT1^PROX^* was almost completely silenced (Figures 4A and 4C). Snf2 was also recruited to the *HNT1* locus within 5 minutes of DTT treatment, and levels of Snf2 binding increased over time in a pattern that strikingly resembled the *HNT1^LUTI^* expression pattern (Figures 4C and 4D), introducing the possibility that Snf2 recruitment is coupled to *HNT1^LUTI^*transcription. Along with this rapid recruitment of the Swi/Snf complex following UPR induction, the -1 and +1 nucleosomes surrounding the *HNT1^PROX^*TSS were also remodeled within 5 minutes of DTT treatment (Figure 4E).

**Figure 4.**
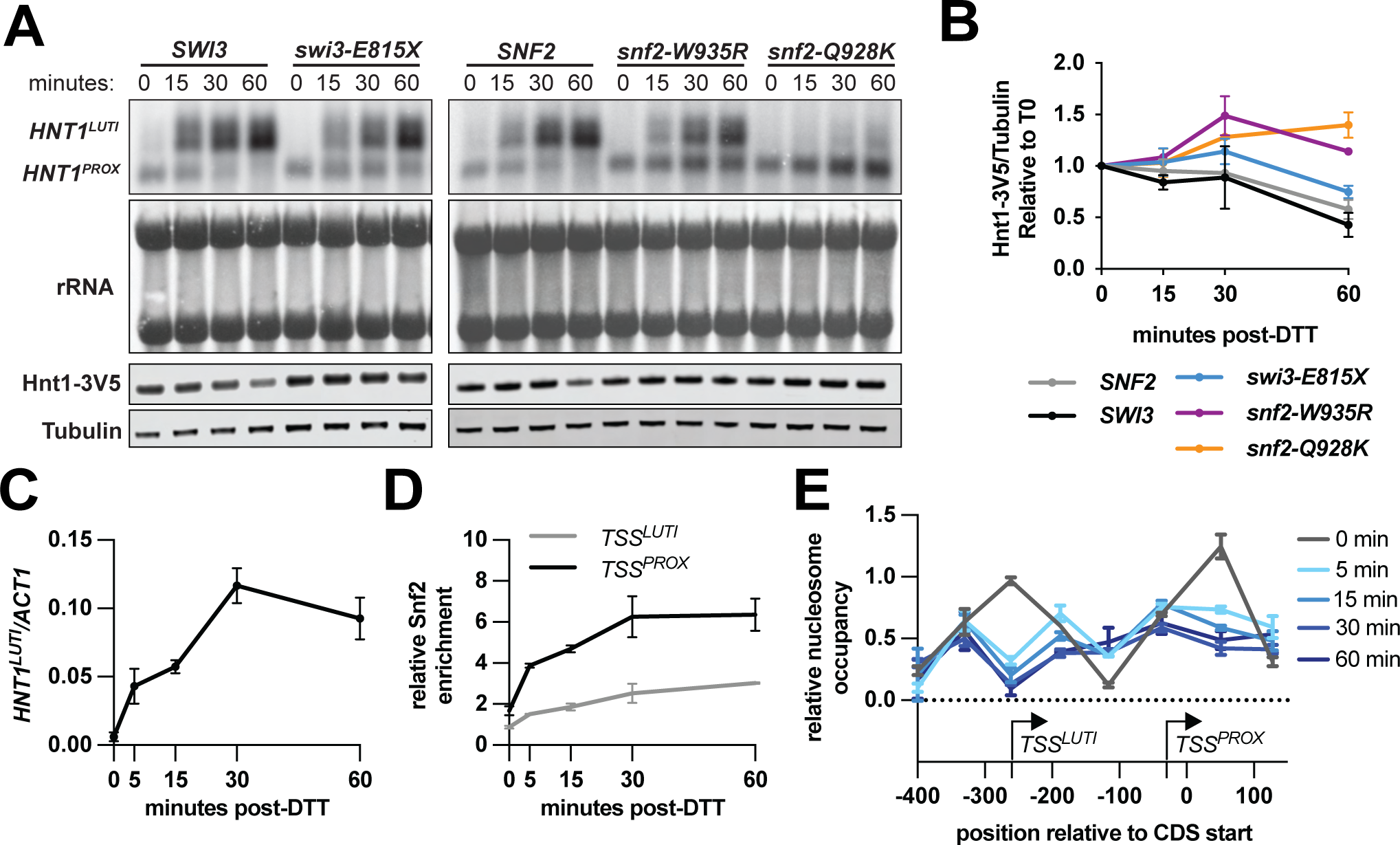
Swi/Snf facilitates rapid and sustained repression of *HNT1^PROX^*upon *HNT1^LUTI^* induction. **A.** RNA blot probed for the *HNT1* CDS (top) and immunoblot against the V5 epitope in strains harboring an *HNT1-3V5* fusion allele (bottom). For the RNA blot, rRNA bands are detected by methylene blue staining. For the immunoblot, alpha-tubulin is used as a loading control. Cells were collected at 0, 15, 30, and 60 minutes after treatment with 5 mM DTT for RNA and protein extraction. Strains: *SWI3* (UB24251), *swi3-E815X* (UB24253), *SNF2* (UB30152), *snf2-W935R* (UB30156), and *snf2-Q928K* (UB30154). One of two biological replicates is shown. **B.** Quantification of immunoblots portrayed in (A) for two biological replicates. **C.** RT-qPCR measuring relative abundance of *HNT1^LUTI^* mRNA in cells (UB29161) collected at 0, 5, 15, 30, and 60 minutes after DTT treatment (n = 2). **D.** Snf2 ChIP-qPCR measuring relative occupancy of Snf2-3V5 (Swi/Snf) at the *HNT1^LUTI^*TSS (black) or *HNT1^PROX^* TSS (gray; n = 2). A primer pair directed against the heterochromatic HMR locus was used as an internal control. Cell collection was done in parallel with cells collected in (C) and (E). **E.** MNase-qPCR measuring relative nucleosome occupancy at the *HNT1* 5′ regulatory region (n = 2). A primer pair directed against the *PHO5* promoter was used as an internal control.

Consistent with the TL-seq results, *snf2-W935R* and *swi3-E815X* cells induced *HNT1^LUTI^* but failed to silence *HNT1^PROX^*, exhibiting higher levels of the *HNT1^PROX^*isoform visible as early as 15 minutes post-DTT treatment (Figure 4A). In contrast, *snf2-Q928K* cells failed to express *HNT1^LUTI^* and exhibited dramatic upregulation of *HNT1^PROX^* (Figure 4A). The high accumulation of *HNT1^PROX^* in this mutant resembled the outcome of deleting the *HNT1^LUTI^* promoter (Figure S4A), suggesting that increased *HNT1^PROX^* expression in *snf2-Q928K* cells was due solely to the lack of *HNT1^LUTI^* expression. Importantly, the *snf2-Q928K* mutant properly activated the UPR response, as indicated by cytosolic splicing of the UPR transcription factor *HAC1* mRNA (Cox and Walter, 1996; Rüegsegger et al., 2001) (Figure S4B), suggesting transcriptional defects in *snf2-Q928K* cells are specific to certain targets and do not result from broadly aberrant stress response. As expected, increased levels of the coding *HNT1^PROX^* mRNA isoform in each of the Swi/Snf LUTI escape mutants resulted in increased Hnt1-3V5 protein levels (Figures 4A and 4B).

### Chromatin changes at the *HNT1* locus depend on transcription initiation and elongation of *HNT1^LUTI^*

We next sought to investigate whether Snf2 recruitment and nucleosome remodeling at the *HNT1* locus is dependent on transcription initiation of *HNT1^LUTI^*, its elongation through downstream chromatin, or both its initiation and elongation. We engineered two LUTI perturbation mutants: one lacking the *HNT1^LUTI^*promoter (*LUTIΔ*) and the other containing an insertion of a strong transcriptional terminator sequence (*CYC1t)* between the *HNT1^LUTI^*and *HNT1^PROX^* TSSs (Figure 5A). Both perturbations eliminated full-length *HNT1^LUTI^* expression (Figure 5B).

**Figure 5.**
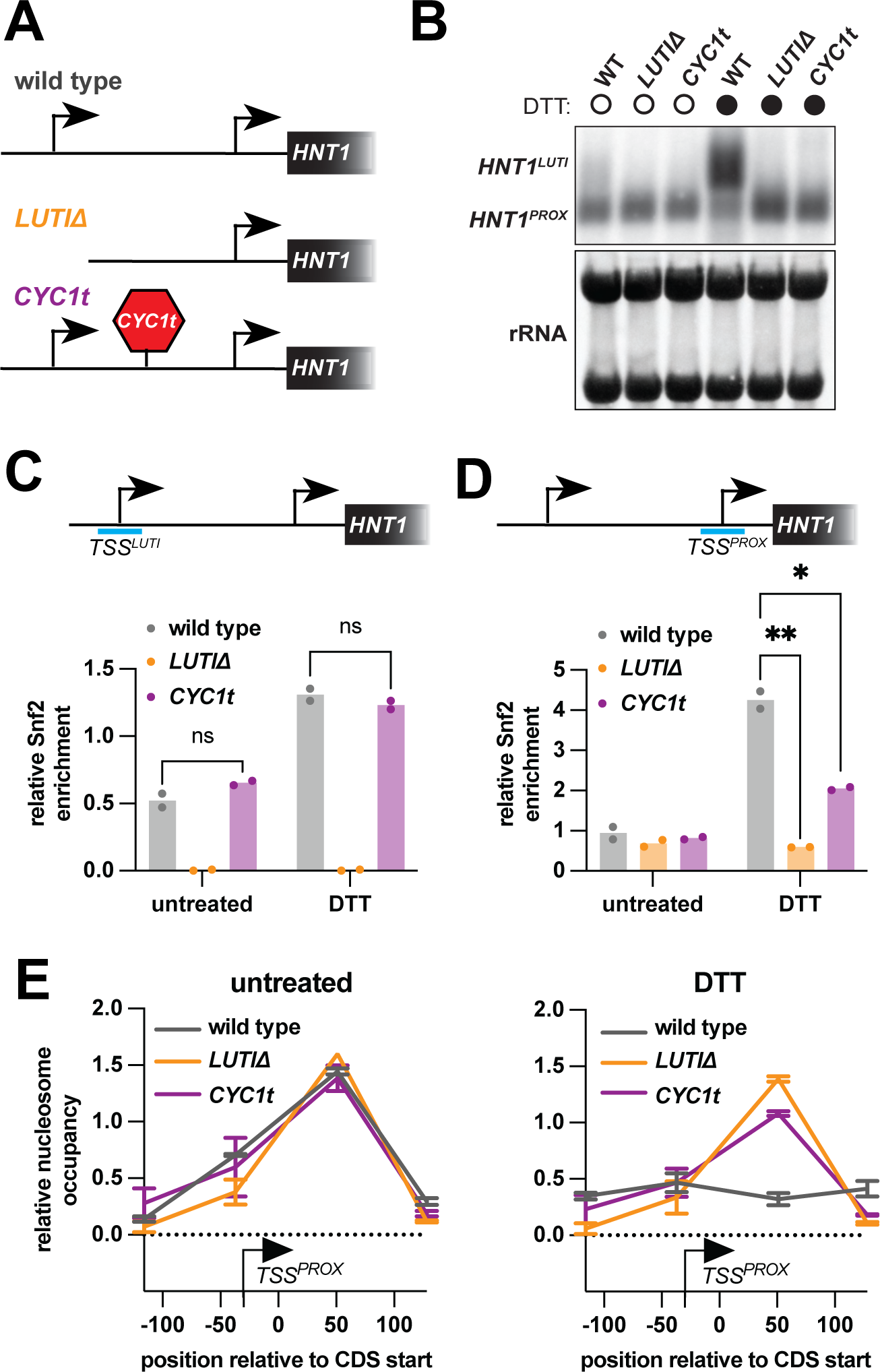
Chromatin changes at the *HNT1* locus depend on transcription initiation and elongation of *HNT1^LUTI^*. **A.** Schematic of *cis* regulatory *HNT1* mutant alleles. **B.** RNA blot probed for the *HNT1* CDS with RNA collected from wild-type (UB32339), *HNT1^LUTIΔ^* (UB32342), and *HNT1^LUTI-CYC1t^* (UB36048) cells that were untreated or treated with 5 mM DTT for 30 min. **C.** Snf2 ChIP-qPCR measuring relative occupancy of Snf2 at the *HNT1^LUTI^* TSS (n = 2). Cells were collected in parallel with cells collected in (B), (D), and (E). **D.** Snf2 ChIP-qPCR measuring relative occupancy of Snf2 at the *HNT1^PROX^* TSS (n = 2). There is significantly decreased Snf2 binding in DTT-treated *HNT^LUTIΔ^* cells (two-way ANOVA, p = 0.0093) and *HNT1^LUTI-CYC1t^* cells (two-way ANOVA, p = 0.0254) compared to wild type. Differences between the *HNT^LUTIΔ^* and *HNT1^LUTI-CYC1t^* cells were not statistically significant (two-way ANOVA, p = 0.0254). **E.** MNase-qPCR measuring relative occupancy for the *HNT1^PROX^* +1 nucleosome.

As the *HNT1^LUTI^* TSS chromatin is deleted in the *LUTIΔ* mutant, the signal for Snf2 binding was reduced to background levels in *LUTIΔ* cells at this position (Figure 5C). Snf2 binding at the *HNT1^LUTI^* TSS was unaffected by *CYC1t* insertion, suggesting *HNT1^LUTI^* initiation is sufficient for Swi/Snf recruitment upon DTT treatment (Figure 5C). Interestingly, binding of Snf2 at the *HNT1^PROX^* TSS was reduced in both *LUTIΔ* and *CYC1t* mutants compared to wild-type cells upon UPR induction (Two-way ANOVA, p = 0.0093 [*LUTIΔ*], p = 0.0254 [*CYC1t*]; Figure 5D). To assay nucleosome remodeling in the LUTI perturbation mutants, we restricted our analysis to positions with shared sequence identity among all three strains, which encompasses the *HNT1^PROX^* +1 nucleosome. In contrast to wild-type cells, this nucleosome was not remodeled in *LUTIΔ* or *CYC1t* cells with UPR induction (Figure 5E).

Altogether, these results revealed that *HNT1^LUTI^* initiation is sufficient for Swi/Snf recruitment to the *HNT1* locus, but downstream Swi/Snf occupancy and nucleosome remodeling at the *HNT1^PROX^* promoter requires productive elongation of *HNT1^LUTI^*. Importantly, the lack of Snf2 binding and nucleosome remodeling at the *HNT1^PROX^* TSS in the *CYC1t* mutant implies remodeling of the *HNT1^PROX^* +1 nucleosome in wild-type cells is an active process by the Swi/Snf complex rather than a result of cascading effects from upstream remodeling at the *HNT1^LUTI^*promoter.

### Swi/Snf regulates gene-body nucleosome occupancy for its canonical gene targets

We wondered whether Swi/Snf-dependent transcriptional interference stems from a general function of the Swi/Snf complex in nucleosome remodeling during transcription elongation. Previous studies have implicated the Swi/Snf complex in transcription elongation (Schwabish and Struhl, 2007; Shivaswamy and Iyer, 2008), however in these cases mutant analysis was performed using *snf2Δ* cells which exhibit severe transcriptional defects, making it difficult to uncouple transcription initiation from elongation phenotypes at target loci. The *swi3-E815X* and *snf2-W935R* mutants, which exhibit few changes in global transcript levels (Figure 2C, Table S3), present a unique opportunity to investigate Swi/Snf regulation of chromatin at its gene targets in a context where transcription levels are largely unperturbed.

First, we generated a list of genes that were significantly downregulated in *snf2Δ* cells compared to wild type under normal, unstressed conditions (DESeq2, adjusted p-value < 0.05). From this list, we next eliminated genes that were not bound by Snf2 based on ChIP-seq data, as these are not direct Swi/Snf targets. Finally, we eliminated genes that were also differentially regulated in either *swi3-E815X* or *snf2-W935R* mutants (Table S3). We did not remove genes affected by the *snf2-Q928K* mutant, as this mutant confers more severe loss of Swi/Snf function, resembling null transcriptional phenotypes for many loci (Figure 2C). This yielded a list of 250 Swi/Snf targets (Figure 6A, left, Table S3). As a control set, we generated a random set of 250 genes whose transcription was not significantly affected by any of the Swi/Snf mutants (DESeq2 adjusted p-value > 0.05) and were not bound by Snf2 (Figure 6A, right, Table S3).

**Figure 6.**
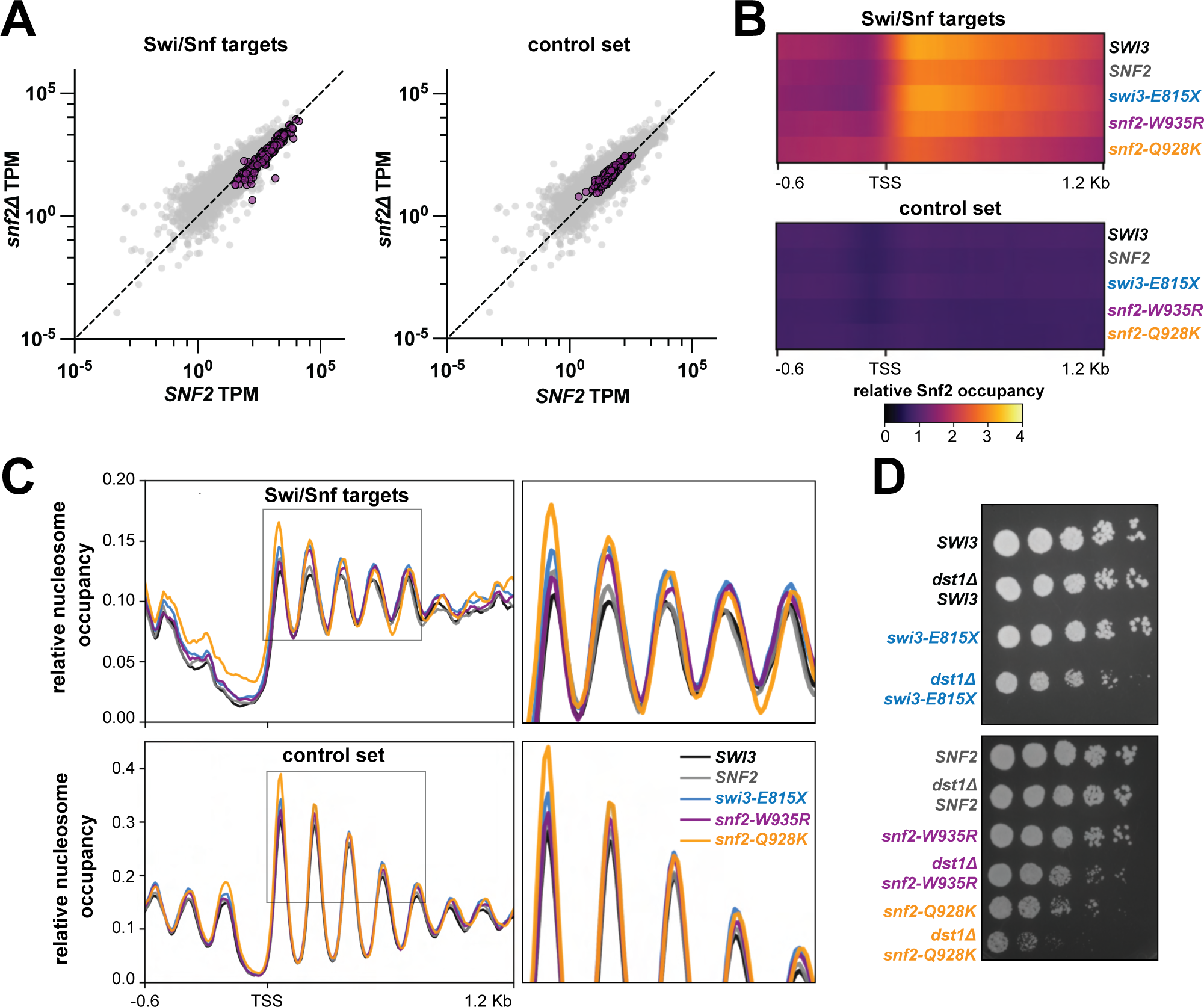
Swi/Snf regulates gene-body nucleosome occupancy for its canonical gene targets. **A.** Scatterplots comparing mRNA abundances (TPM) between *snf2Δ* (UB29781) and *SNF2* (UB28914) cells. Genes that are Swi/Snf-regulated (left, n = 250) or genes from a non-regulated control set (right, n = 250) are highlighted in purple. **B.** Heatmaps portraying normalized Snf2 occupancy generated from Snf2 ChIP-seq data for genes that are Swi/Snf-regulated (top) or the non-regulated control set (bottom). Strains: *SWI3* (UB30070), *SNF2* (UB30387), *swi3-E815X* (UB30071), *snf2-W935R* (UB30391), and *snf2-Q928K* (UB30389). **C.** Metagene plots created from MNase-seq data for genes that are Swi/Snf regulated (top) or the non-regulated control set (bottom). Strains are the same as in (B). **D.** Serial dilution and plating growth assay on YPD media. Strains: *SWI3* (UB19205), *SWI3 dst1Δ* (UB36182), *swi3-E815X* (UB19209), *swi3-E815X dst1Δ* (UB28096), *SNF2* (UB28914), *SNF2 dst1Δ* (UB36185), *snf2-W935R* (UB28922), *snf2-W935R dst1Δ* (UB36186), *snf2-Q928K* (UB28915), and *snf2-Q928K dst1Δ* (UB36188).

We next compared average Swi/Snf occupancy and nucleosome profiles for these two sets of genes between wild type and LUTI escape mutants. Importantly, Snf2 ChIP-Seq and MNase-Seq experiments were performed with spike-in control cells from a divergent strain of *S. cerevisiae*, enabling us to compare between samples after scaling read coverage with the calculated normalization factor (Vale-Silva et al., 2019) (Table S3). As expected, LUTI escape mutants did not exhibit Snf2 binding or nucleosome profile differences for the control set of genes, apart from increased occupancy of the -1 and +1 nucleosome in *snf2-Q928K* cells that may be due to the higher degree of pleiotropy in this mutant (Figures 6B and 6C). The *swi3-E815X* and *snf2-W935R* mutants exhibited normal binding of Snf2 for Swi/Snf targets, with average Snf2 occupancy peaking at the region encompassing +1, +2, and +3 nucleosomes (Figures 6B and 6C). The *snf2-Q928K* mutant, however, exhibited slightly lower Snf2 occupancy at positions downstream of the TSS (Figure 6B). On average, the *snf2-Q928K* mutant also exhibited increased nucleosome occupancy within the NDR and at downstream nucleosomes for Swi/Snf target genes (Figures 6C and S6C-D). Interestingly, the *swi3-E815X* and *snf2-W935R* mutants did not impact the chromatin at the TSS or +1 nucleosome as strongly as the *snf2-Q928K* mutant but did exhibit increased nucleosome occupancy for gene-body nucleosomes, especially for the +2 nucleosome (Figure 6C and S6C-D).

Given their effects on gene-body nucleosomes, we wondered whether the *swi3-E815X* and *snf2-W935R* mutations confer specific defects in co-transcriptional nucleosome remodeling for Swi/Snf targets. While these mutants do not impair transcription elongation to a degree that impacts transcript levels (Figures S6A and S6B), it is possible that the activity of other transcription elongation factors compensates for loss of Swi/Snf function to promote normal elongation by Pol II. To address this possibility, we assayed synthetic phenotypes for LUTI escape mutants combined with deletion of *DST1,* a gene encoding the general elongation factor TFIIS which alleviates Pol II stalling and is commonly synthetic lethal with deletion of other elongation factors (Costa and Arndt, 2000; Davie and Kane, 2000; Malagon et al., 2004). Consistent with previous observations that *snf2Δ dst1Δ* cells are inviable (Davie and Kane, 2000; Schwabish and Struhl, 2007), the *dst1Δ* combined with the *snf2-Q928K* mutation resulted in a profound growth defect (Figure 6D). The *swi3-E815X* and *snf2-W935R* mutants also displayed growth defects when combined with *dst1Δ* (Figure 6D), in contrast to their minimal effects on Swi/Snf loss-of-function phenotypes that are dependent on Swi/Snf’s gene activation function (Figures 2B and S2B-C). From these results, we concluded that the *swi3-E815X* and *snf2-W935R* mutants confer broad defects in co-transcriptional remodeling. Furthermore, these LUTI escape mutants allow for canonical Swi/Snf function in transcriptional activation to be uncoupled from an understudied role for Swi/Snf in transcription elongation.

## DISCUSSION

The Swi/Snf complex is a highly conserved chromatin remodeler that has been extensively studied for its role in transcriptional activation. Mutations in Swi/Snf subunits are found in ∼20% of all human tumors, making the complex one of the most commonly affected in cancer (Kadoch and Crabtree, 2015). Interestingly, a few studies have provided evidence in support of Swi/Snf acting in gene repression (Choi et al., 2015; Martens and Winston, 2002; Menon et al., 2019; Murphy et al., 1999; Zhu et al., 2013), however, it has remained unclear whether Swi/Snf-dependent transcriptional repression occurs directly or indirectly as a result of transcription initiation defects at nearby loci or decreased expression of other transcriptional regulators.

In this study, we have demonstrated a direct role of the Swi/Snf complex in transcriptional repression. This occurs by co-transcriptional nucleosome remodeling by Swi/Snf downstream of distal TSSs expressing non-canonical transcripts including LUTIs. Consequently, the nucleosome depleted region of the CDS-proximal promoter, which resides downstream of the distal TSS becomes nucleosome occupied, thereby resulting in the repression of the protein-coding transcript isoform (Figure 7, top). LUTI escape mutants *swi3-E815X* and *snf2-W935R* have minimal effects on Swi/Snf’s ability to facilitate transcription initiation at its target loci. Instead, these mutations specifically disrupt nucleosome remodeling at positions downstream of the active TSS, resulting in the repression of the TSS^PROX^ for Swi/Snf’s TSS^DIST^ targets (Figure 7, middle). With more severe loss of Swi/Snf function, as in null or *snf2-Q928K* mutants, remodeling at the active TSS is reduced, leading to impaired transcription initiation. In these cases, transcriptional interference at the TSS^PROX^ for Swi/Snf’s TSS^DIST^ targets is reduced indirectly, as a result of lower TSS^DIST^ transcription (Figure 7, bottom).

**Figure 7.**
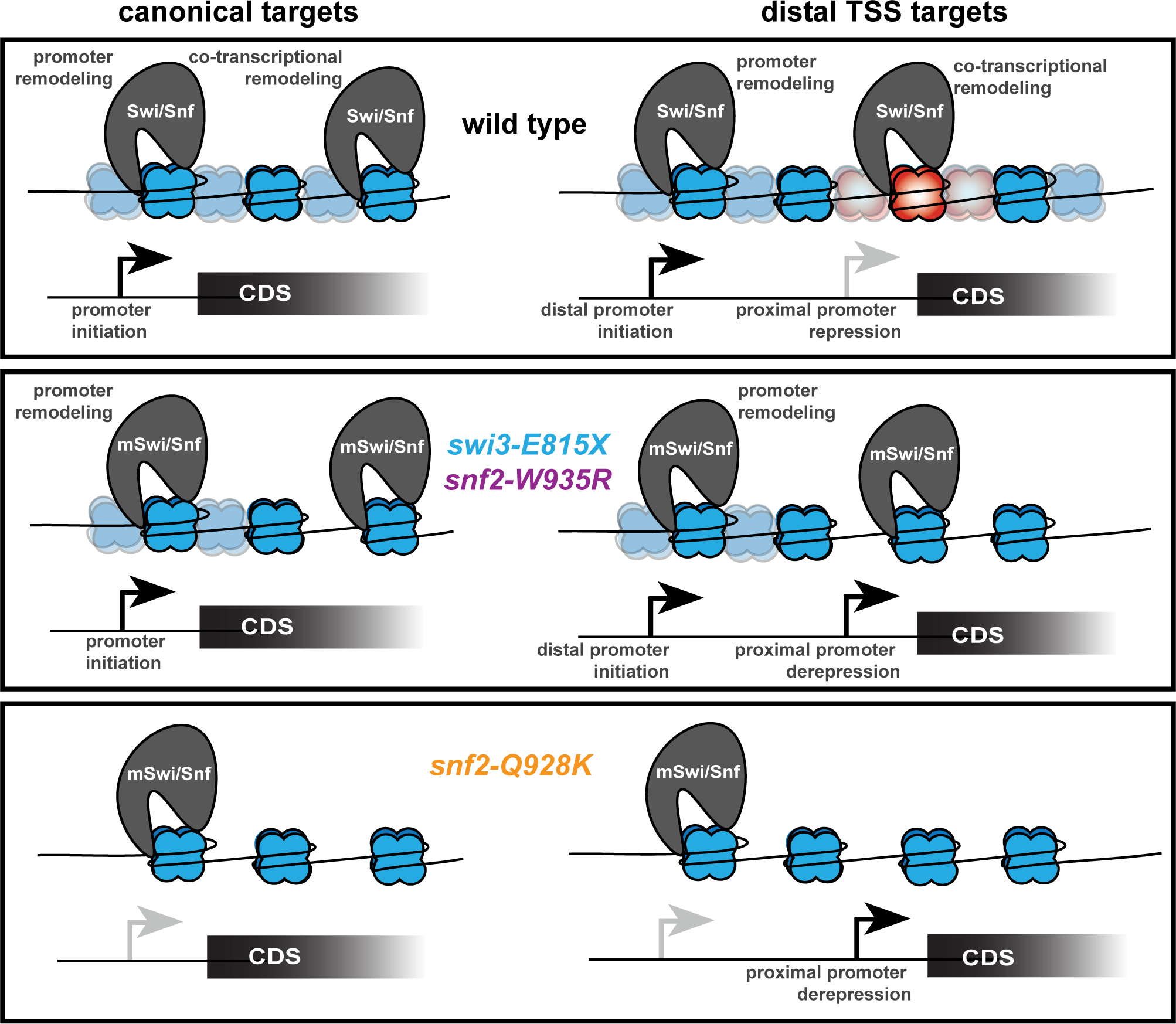
Model for Swi/Snf regulation of TSS activation and repression. In wild-type cells (top), the Swi/Snf complex is recruited to canonical promoters (left) or distal promoters (right) and performs nucleosome remodeling to aid in transcription initiation. Swi/Snf also performs a secondary function at its targets to remodel nucleosomes downstream of the active TSS, which represses TSS^PROX^ promoters for its TSS^DIST^ targets. Productive remodeling is indicated with translucent, “fuzzy” nucleosomes and transcriptionally interfering nucleosomes are indicated in red. Middle: in *swi3-E815X* and *snf2-W935R* mutants, transcription initiation function by the Swi/Snf complex remains intact but downstream remodeling is impaired. For canonical targets, these mutants compromise nucleosome remodeling within the gene body without compromising transcription levels. For TSS^DIST^ targets, reduced nucleosome remodeling downstream of the active TSS results in derepression of the TSS^PROX^. Bottom: in *snf2Q928K* cells, nucleosome remodeling and transcription initiation at both canonical and TSS^DIST^ promoters is reduced. Reduced transcriptional readthrough results in derepression of the _TSSPROX._

### Phenotypic differences among LUTI escape mutants

Our in-depth characterization uncovered striking phenotypic differences between the two missense mutations within the helicase domain of Snf2, *snf2-W935R* and *snf2-Q928K*. Binding of Snf2 is not reduced at Swi/Snf target loci in either mutant (Figures S3A and 6B), indicating these mutations confer remodeling defects rather than recruitment defects. How might these LUTI escape mutations impact Swi/Snf remodeling function? Mapping these conserved residues on the cryo-EM structure of the Swi/Snf complex bound to a nucleosome revealed that the Q928 residue of Snf2 directly contacts nucleosomal DNA, whereas the W935 residue resides in a nearby pocket of Snf2 that does not directly contact DNA or histones (Figure 1E). This structural information, combined with our findings that the *snf2-Q928K* impairs Swi/Snf function more severely than *snf2-W935R*, suggests the glutamine residue at position 928 is critical for Snf2 remodeling function. Several cancer-associated mutations in humans also affect residues in the nucleosome binding region of the *SNF2* homolog *BRG1* (K. Chen et al., 2023), indicating this binding interface is functionally conserved.

Current models for Swi/Snf nucleosome remodeling propose ATP hydrolysis by Snf2 promotes DNA translocation and nucleosome sliding (Bowman, 2010; K. Chen et al., 2023; M. Li et al., 2019). Both the *snf2-W935R* and *snf2-Q928K* mutations result in a neutral-to-positive charge substitution, which may impair Snf2’s ability to translocate nucleosomal DNA due to an increased affinity for the negatively charged DNA backbone. Another possibility is that missense mutations at nucleosome-binding sites in Snf2 lead to structural changes that impair its ATPase activity. In support of this notion, a *snf2-W935A* mutant was previously shown to reduce ATPase activity to 80% that of wild-type levels (Smith and Peterson, 2005). Although the Q928 residue has not been interrogated for a role in ATPase function, Snf2 mutations that eliminate ATPase activity result in dominant negative phenotypes (Richmond and Peterson, 1996), indicating the dominant negative phenotype for the *snf2-Q928K* allele uncovered in this study may stem from defects in ATP binding or hydrolysis.

The *swi3-E815X* mutation results in a truncation of Swi3 at its C-terminal coiled-coil domain. As Swi3 is thought to act as a scaffold for complex assembly (Han et al., 2020; Yang et al., 2007), defects in Swi/Snf remodeling could arise from structural changes in this mutant or reduced interactions between the complex with other transcription factors. In the case of the *swi3-E815X* and *snf2-W935R* mutants, which primarily disrupt Swi/Snf co-transcriptional remodeling without other pleiotropic defects, it is possible the mutations confer structural changes that inhibit key interactions between the Swi/Snf complex and elongation factors. Additional work to investigate structural changes induced by LUTI escape mutations and physical interactions between Swi/Snf subunits with transcription elongation factors is required to assess how the LUTI escape mutations differentially impact Swi/Snf function in transcription initiation and elongation.

### One of several transcriptional interference mechanisms: disrupting promoter architecture

Eukaryotic promoters consist of a nucleosome free region (NFR) or nucleosome depleted region (NDR) flanked by two well-positioned nucleosomes with acetylated histones (Bai and Morozov, 2010; Venkatesh and Workman, 2015). While the TSS for most genes lies within 10-15 base pairs of the 5’ end of the +1 nucleosome, the NFR or NDR is thought to allow access for sequence-specific transcription factors and the pre-initiation complex to start transcription at the gene promoter (Gill et al., 2020; Jansen and Verstrepen, 2011; Rando and Winston, 2012). In the case of LUTIs and other transcripts with distal TSSs relative to the CDS-proximal TSS (TSS^PROX^), transcription proceeds across the TSS^PROX^, subjecting the proximal promoter to co-transcriptional chromatin changes that normally function to promote elongation and inhibit cryptic transcription initiation. Several transcriptional interference pathways have already been uncovered, many of which involving histone modification or nucleosome remodeling (Carrozza et al., 2005; Hartzog et al., 1998; Hennig et al., 2012; Pruneski et al., 2011; Smolle et al., 2012). For Swi/Snf-repressed TSS^PROX^ loci uncovered in this study, it seems that Swi/Snf remodeling of the -1 and +1 nucleosomes surrounding the proximal promoter contributes to TSS^PROX^ repression, possibly by creating increased nucleosome mobility into what was previously the NFR/NDR.

Based on this study along with previous findings, Swi/Snf remodeling of nucleosomes downstream of the TSS^DIST^ is one of several possible routes to achieve transcriptional interference of the TSS^PROX^. There may, in fact, be other loci at which Swi/Snf acts in parallel with other described transcriptional interference pathways during UPR induction that evaded our detection due to compensation by other factors in the LUTI escape mutants. Based on our observations that the *snf2-Q928K* mutant impairs transcription of the distal isoform (Figure 3F) and that activation of *HNT1^LUTI^* is sufficient for Snf2 recruitment (Figure 5C), it seems that recruitment of the Swi/Snf complex to the distal promoter is a prerequisite for the downstream transcriptional interference activity by the complex. Recruitment of Swi/Snf may be mediated through interactions with acetylated histones or sequence-specific transcription factors at distal promoters. In support of this notion, Swi/Snf occupancy at several canonical UPR-induced promoters depends on the master UPR transcription factor Hac1 (Sahu et al., 2021). Swi/Snf interaction with Hac1 may also be the basis for its recruitment to *HNT1^LUTI^*, a reported Hac1 target (Van Dalfsen et al., 2018). Another prerequisite for Swi/Snf-dependent transcriptional interference may be shorter distance between the two TSSs, based on our observation that *swi3-E815X* and *snf2-W935R* do not strongly affect nucleosomes further downstream from the TSS (Figures 6C and S6D) and previous observations that Snf2 is most highly enriched at the -1, +1, +2, and +3 nucleosomes for its gene targets (Yen et al., 2012). Genes with greater distances between the two TSSs may rely on other mechanisms for transcriptional interference to occur, such as the H3K36me3 pathway (J. H. Kim et al., 2016).

### Roles for Swi/Snf beyond transcriptional activation

While most investigations into Swi/Snf cellular function have focused on its role as a co-activator at gene promoters, several studies have uncovered evidence indicating the Swi/Snf complex also functions in transcription elongation. For example, Snf2 binds with similar patterns and kinetics as Pol II along coding regions upon induction of transcription (Schwabish and Struhl, 2007). In particular, Snf2 occupies promoters and coding regions for heat-shock induced genes (Shivaswamy and Iyer, 2008) and transcription elongation of human *HSF1* is necessary for Swi/Snf recruitment at the *HSF1* locus (Corey et al., 2003). Finally, *snf2Δ* cells exhibit increased nucleosome occupancy at promoters and within gene bodies (Rawal et al., 2018). However, these studies utilize a *snf2Δ* null mutant, which reduces transcription levels at Swi/Snf target loci. In *snf2Δ* cells, it is difficult to distinguish whether differences in nucleosome occupancy within the gene body are due to reduced transcription versus loss of Swi/Snf activity on gene-body nucleosomes and what the biological significance of Swi/Snf-mediated co-transcriptional remodeling is. Here, we provide conclusive evidence that the Swi/Snf complex functions in co-transcriptional nucleosome remodeling by uncovering specific mutants and genomic contexts that impair nucleosome remodeling within gene bodies without reducing transcript levels. Although overall transcript levels are not impaired for most Swi/Snf targets in the *swi3-E815X* and *snf2-W935R* mutants, we cannot eliminate the possibility that these mutations affect transcription elongation rates or Pol II stalling frequency. Future work to specifically assay transcription elongation would aid in determining whether the phenotypes associated with LUTI escape mutants stem from altered transcription elongation rates.

Previous studies have also uncovered a role for the Swi/Snf complex in gene repression in yeast as well as more complex organisms (Choi et al., 2015; Martens and Winston, 2002; Menon et al., 2019; Murphy et al., 1999; Zhu et al., 2013). In yeast, the Swi/Snf complex activates transcription of the non-coding transcript *SRG1* in serine replete conditions. *SRG1* transcription results in repression of the *SER3* promoter via co-transcriptional nucleosome deposition by the Paf1 complex, Spt6, Spt16, and Spt2 (Martens et al., 2005; Martens and Winston, 2002; Pruneski et al., 2011). In this case, Swi/Snf-based repression of *SER3* is indirect, whereas direct repression at the *SER3* promoter is achieved by other elongation factors, exemplifying the complexity and diversity among transcriptional interference pathways.

Genome-wide studies in yeast have revealed that deletion of Swi/Snf genes results in both downregulation and upregulation of transcription for subsets of genes (Dutta et al., 2017; Sen et al., 2017). While upregulation of a gene in a *snf2Δ* mutant may indicate the Swi/Snf complex normally represses its transcription, it is difficult to conclude whether repression is achieved through Swi/Snf remodeling at that locus or indirect effects given the degree of pleiotropy in *snf2Δ* cells. By integrating transcriptome, genomic Snf2 occupancy, and genomic nucleosome data for wild-type cells compared to LUTI escape mutants in this study, we were able to infer with high confidence that the Swi/Snf complex induces bona fide TSS^DIST^ targets and represses transcription at downstream TSS^PROX^ promoters via co-transcriptional nucleosome remodeling.

### Concluding remarks

In summary, we uncovered a direct role for the Swi/Snf complex in transcriptional interference using genetics to reveal novel regulators of LUTI-based gene repression. In line with its characterized role in gene activation, the Swi/Snf complex activates transcription of non-canonical mRNAs initiating from CDS-distal promoters in response to protein folding stress, including three LUTI mRNAs. We found that the Swi/Snf complex also performs a secondary function at these loci to repress the TSS^PROX^ via nucleosome remodeling downstream of the TSS^DIST^. For the LUTI-regulated gene *HNT1*, proper Snf2 occupancy, nucleosome remodeling, and *HNT1^PROX^*repression were dependent on *HNT1^LUTI^* transcriptional initiation and elongation, indicating repression of *HNT1^PROX^* is achieved via co-transcriptional nucleosome remodeling by Swi/Snf.

Our finding that the Swi/Snf complex can act as both a transcriptional activator and repressor simultaneously at the same genomic locus is relevant to ongoing investigations into the link between Swi/Snf mutations and cancer. The transcriptional landscape in human cells is complex, with a high prevalence of non-coding transcription and transcript isoform toggling (Hangauer et al., 2013; Wang et al., 2016). It is thought that Swi/Snf mutations lead to disease through both downregulation and upregulation of gene expression (Cenik & Shilatifard, 2021; Lu and Allis, 2017), however mechanisms for Swi/Snf-based gene repression have remained unclear. Our findings indicate one route through which Swi/Snf chromatin remodeling can lead to direct gene repression in yeast: transcriptional interference of coding transcripts. Thus, future work to investigate whether human Swi/Snf complexes also play a role in transcriptional interference for genes with tandem promoters, such as the LUTI-regulated proto-oncogene *MDM2* (Hollerer et al., 2019), will reveal whether transcriptional interference activity by the Swi/Snf complex is evolutionarily conserved.

## MATERIALS AND METHODS

### Yeast strains

All yeast strains were derived from the W303 background. Genotypes are listed in Table S4. All gene deletions were made via Pringle-based insertion of a drug resistance cassette replacing the endogenous CDS, except for the case of the *hnt1Δ* in which the CDS and 275 bases upstream of the CDS was replaced with a drug resistance cassette (Bähler et al., 1998). The *pLexO-HIS3^LUTI^* reporter construct was generated by cloning the *NDC80^LUTI^* 5′ leader sequence and the *HIS3* CDS into a *LEU2* single integration vector harboring a 3X *lexO* array upstream of a *CYC1* minimal promoter via three-piece Gibson Assembly (Gibson et al., 2009). The *pTEF1-ADE2^LUTI^* reporter construct was derived from the *pLexO-HIS3^LUTI^* in a two-step process: First the *HIS3* CDS was replaced with the *ADE2* CDS by two-piece Gibson Assembly to create a *pLexO-ADE2^LUTI^* plasmid, then the Lex-inducible promoter was replaced with the *TEF1* promoter amplified from gDNA by two-piece Gibson Assembly. Transgenic *SNF2* and *SWI3* wild-type or LUTI escape alleles with their endogenous gene promoter and terminator sequences were amplified from genomic DNA and cloned into a *LEU2* single integration plasmid via two-piece Gibson Assembly.

For strains used in Snf2 ChIP experiments, Pringle-based insertion was used to insert a 3V5 epitope tag at the C-terminal end of the endogenous *SNF2* CDS (for UB29161, UB30070, and UB30071), or the *LEU2* transgenic copy of *SNF2, snf2-W935R,* and *snf2-Q928K* (for UB30387, UB30391, and UB30389). Wild type and *pDISTΔ* transgenic constructs for *HNT1*, *ADI1*, *ODC2*, and *ERG27* were generated by cloning the CDS for each gene with upstream sequence including the distal TSS and 300 bases upstream (WT) or the sequence immediately downstream of the the distal TSS (*pDISTΔ*) into a *TRP1* single integration vector harboring a C-terminal 3V5 epitope sequence. The *HNT1^LUTI-CYC1t^* construct was generated by cloning the *CYC1* terminator into the *HNT1^LUTI^-3V5 TRP1* single integration vector 40 bp downstream of the *HNT1^LUTI^* TSS via two-piece Gibson Assembly. For all Gibson Assembly reactions, the NEBuilder HiFi Master Mix was used according to kit instructions (E5520S, *New England Biolabs*). All single integration plasmids were digested with PmeI before transformations and integrations were verified by PCR. All strains and plasmids used in this study are available upon request.

### Growth conditions

For selecting LUTI escape mutants, cells were grown overnight at 30°C in liquid YPD (1% yeast extract, 2% peptone, 2% dextrose, tryptophan (96 mg/L), uracil (24 mg/L), adenine (12 mg/L) to saturation. A final concentration of 25 nM β-estradiol was added to overnight YPD flasks to deplete intracellular His3 protein. The day of plating, cells were back-diluted to an OD_600_ = 0.1, then grown at 30°C until they reached OD_600_ ∼0.6. After quantifying cell density using a hemacytometer, cells were pelleted by microcentrifugation (2 min at 2000 g), washed with sterile water, and plated on synthetic complete media (SC; 6.7 g/L yeast nitrogen base without amino acids, 2% dextrose, 20 mg/L adenine, 20 mg/L arginine, 60 mg/mL leucine, 20 mg/mL tryptophan, 20 mg/L methionine, 30 mg/L lysine, 30 mg/L tyrosine, 50 mg/L phenylalanine, and 200 mg/L threonine) lacking histidine with 25 nM β-estradiol and 200 µM 3-amino-1,2,4-triazole (3-AT) to completely silence His3 activity in control cells. Cells were plated at a density of 3 million cells/25 ml plate and grown for four days at 30°C. Viable colonies were selected after three and four days post-plating.

For serial dilution and plating assays, cells were collected from an overnight YPD plate grown at 30°C and resuspended in water to a concentration of OD_600_ = 0.2. Next, four 1:5 serial dilutions were performed and 2.5 ul of each dilution was plated. For *HIS3^LUTI^* phenotyping, cells were plated on a control plate (SC -his with 200 µM 3-AT) and a selective plate (SC -his with 200 µM 3-AT and 25 nM β-estradiol) then allowed to grow for three days at 30°C prior to imaging. For *dst1Δ* phenotyping, cell dilutions were plated on YPD then allowed to grow for two days at 30°C prior to imaging. *ADE2^LUTI^* phenotyping was performed by streaking cells grown overnight at 30°C on a YPD plate to a YPD plate lacking supplemental adenine. Cells were then grown overnight at 30°C and imaged. Cells for growth curve experiments were grown overnight at 30°C in liquid YPD then back-diluted to an OD_600_ = 0.1 in liquid YPD or YPS (YPD recipe with 2% sucrose instead of 2% dextrose) in a 96-well plate. Cells were transferred to a 96-well plate in triplicate, sealed with a Breathe Easy Cover (*Sigma-Aldrich*), and grown at 30°C for 24 hours in a plate reader. Absorbance readings were collected every 15 minutes at absorbance = 600 nm with agitation before each reading.

For all cell collections, yeast cells were grown in liquid YPD. Cells were grown at 30°C overnight prior to collection, back-diluted to OD_600_ = 0.1 the day of collection and grown to mid-log phase at 30°C (OD_600_ ∼0.6 for RNA and protein collections) or late-log phase (OD_600_ ∼0.8 for ChIP and MNase collections). For UPR-induction, dithiothreitol (DTT) was added to liquid cultures to a final concentration of 5 mM when cells reached mid- or late-log phase. For all DTT sequencing experiments, cells were split at mid-log (TL-seq) or late-log (ChIP-seq and MNase-seq) phase into a 5 mM DTT-treated flask or untreated control flask and harvested 1 hour post-induction. For DTT time course experiments, cells were collected at 0 min once they had reached mid- or late-log phase, UPR stress was induced with 5 mM DTT, and cells were collected at indicated time points following induction. For *HNT1^LUTI-CYC1t^* and *HNT1^lutiΔ^* experiments, cells were split at late-log phase into a 5 mM DTT-treated flask or untreated control flask and harvested 30 min post-induction.

### DNA extractions for sequencing

Cells were grown in 2 ml liquid YPD cultures overnight at 30°C. 1.5 ml of culture was pelleted by microcentrifugation, supernatant was removed and cells were lysed with ∼300 mg 500 micron acid-washed glass beads, 500 μl lysis buffer (2% Triton X-100, 1% SDS, 100 mM NaCl, 10 mM Tris-HCl pH 8, 1 mM EDTA), and 500 μl 25:24:1 phenol:chloroform:isoamyl alcohol. Cells were vortexed in the lysis mixture for 5 min at room temperature and the aqueous phase was separated by microcentrifugation (5 min at 20,000 g). A second extraction was performed in 25:24:1 phenol:chloroform:isoamyl alcohol and DNA was precipitated in 1 ml 100% ethanol at room temperature. The pellet was resuspended in 400 μl TE buffer (10 mM Tris-HCl, 1 mM EDTA, pH 8) and treated with 30 μg RNase A for 30 min at 37°C. The RNase-treated DNA was precipitated in 1 ml 100% ethanol with 10 μl 4 M ammonium acetate and pelleted by microcentrifugation (2 min at 20,000 g). After decanting the pellets were air-dried and resuspended in 50 μl TE. DNA concentration was measured using the Qubit DNA BR Assay Kit (Q32853, *ThermoFisher Scientific*).

### RNA extraction for mRNA-seq, TL-seq, and direct mRNA-seq

Cells were collected by vacuum filtration and snap frozen in liquid nitrogen. For mRNA-seq, ∼15 OD_600_ units of cells were collected. For TL-seq and direct mRNA-seq, ∼100 OD_600_ units of cells were collected. Cells were thawed on ice and resuspended in 400 μl TES buffer (10 mM Tris pH 7.5, 10 mM EDTA, 0.5% SDS) per 15 OD_600_ units. An equal volume of citrate-buffered acid phenol (pH 4.3, P4682, *Sigma-Aldrich*) was added to cells, and they were incubated at 65°C for 30 minutes in a Thermomixer C (*Eppendorf*) shaking at 1400 RPM. After microcentrifugation (20,000 g for 10 min) the aqueous phase was transferred to a second tube with 350 μl chloroform. The aqueous phase was separated by microcentrifugation (20,000 g for 5 min) and RNA was precipitated in 100% isopropanol with 350 mM sodium acetate (pH 5.2) overnight at –20°C. Pellets were washed with 80% ethanol and resuspended in DEPC water for 10 min at 37°C. Total RNA was quantified using the Qubit RNA BR Assay Kit (Q10211, *ThermoFisher Scientific*).

### mRNA sequencing (mRNA-seq)

RNA-seq libraries for three biological replicates were generated with the NEXTflex^TM^ Rapid Directional mRNA-Seq Bundle with poly(A) beads (NOVA-5138, *Bioo Scientific*) according to manufacturer’s instructions. 10 µg of total RNA was used as input for all libraries. AMPure XP beads (A63881, Beckman Coulter) were used to select fragments between 200-500 bp. Libraries were quantified using the Agilent 4200 TapeStation (*Agilent Technologies, Inc*). Samples were submitted for 100 bp SE sequencing by the Vincent J. Coates Genomics Sequencing Laboratory with a NovaSeq SP.

### Poly(A) selection for TL-Seq and direct mRNA-seq

Poly(A) selection was performed on 600 μg of RNA using the Poly(A)Purist^TM^ MAG kit (AM1922, *Ambion*) following manufacturer’s instructions. Poly(A) RNA was precipitated at -20°C for 18 hours in 1.1 ml 100% ethanol with 40 μl 3 M sodium acetate and 1 μl glycogen (5 mg/ml), washed with 1 ml 80% ethanol, dissolved in 21 μl nuclease-free water, and quantified with a Qubit using the RNA BR assay kit (Q10211, *ThermoFisher Scientific*).

### Transcript Leader Sequencing (TL-Seq)

TL-seq was performed on two biological replicates for each sample as described in (Tresenrider et al., 2022) with minor modifications. 5-15 μg poly(A)-selected mRNA was fragmented for 3 minutes at 70°C using alkaline hydrolysis fragmentation reagent (AM8740, *Ambion*). Fragmented mRNAs were purified by performing a 1.8X bead cleanup with RNAClean XP beads (A63987, *Beckman Coulter*). Fragments were dephosphorylated with 20 units Shrimp Alkaline Phosphatase (rSAP, M0371, *New England Biolabs*) for 1 hour at 37°C with 1 μl RNaseOUT (10777019, *Invitrogen*), then purified by acid phenol extraction at 65°C for 45 min in a thermomixer with shaking at 1400 RPM. Dephosphorylated mRNA fragments were precipitated in 100% ethanol with 40 μl 3 M sodium acetate and 1 μl linear acrylamide (AM9520, *Ambion*) at -20°C for >16 hours, washed with 80% ethanol, and resuspended in nuclease-free water. Samples were then treated with 5 units Cap-Clip acid pyrophosphatase (C-CC15011H, *Tebu-Bio*) with 1 μl RNaseOUT. To control for inefficient phosphatase activity in the previous step, a sample from wild-type cells in each condition (untreated or 5 mM DTT) that was not subject to decapping was carried forward along with the Cap-Clip treated samples. Samples were once again purified by acid phenol extraction and ethanol precipitation, then ligated to a custom 5′ adapter (10 μM oligocap) by T4 RNA Ligase I (M0437, *New England Biolabs*) with 1 μl RNaseOUT for 16 hours at 16°C. The ligation reaction was purified by performing a 1.8X bead cleanup with RNAClean XP beads and eluted in 12 μl nuclease-free water. Purified 5′ ligated RNAs were mixed with 1 μl random hexamers (50 μM, N8080127, *ThermoFisher Scientific*), 1 μl dNTP mix (10 mM, 18427013, *Invitrogen*), and 1 μl RNaseOUT then denatured at 65°C for 5 min and cooled on ice. Reverse transcription reactions were performed using SuperScript IV reverse transcriptase (18090010, *Invitrogen*). RNA templates were then degraded by incubating reactions with 5 units of RNase H (M0297, *New England Biolabs*) and 1 μl RNase cocktail enzyme mix (AM2296, *Ambion*). cDNA products were purified by performing a 1.8X bead cleanup with AMPure XP beads (A63881, *Beckman Coulter*), eluted in 23.5 μl nuclease-free water, then subject to second-strand synthesis using 1.5 μl biotinylated oligo (10 μM) and 25 μl KAPA Hi-Fi hot start ready mix 2x (KK2601, *Roche*). Second strand reactions were incubated at 95°C for 3 minutes, 98°C for 15 s, 50°C for 2 minutes, 65°C for 15 minutes and held at 4°C. Double stranded DNA (dsDNA) was purified by performing a 1.8X AMPure XP bead cleanup and quantified using the Qubit dsDNA HS assay kit (Q32851, *Invitrogen*).

All dsDNA (ranging from 2-12 ng per sample) was carried forward into end repair, adenylation, and adapter ligation using the NEXTflex^TM^ Rapid DNA-Seq Kit 2.0 Bundle (NOVA-5188-12, *Bioo Scientific*) according to manufacturer’s instructions. Following post-ligation cleanup and prior to PCR amplification, samples were bound to MyOne Streptavidin C1 Dynabeads^TM^ (65001, *ThermoFisher Scientific*) to capture biotinylated dsDNA. Library amplification over 14-15 PCR cycles, depending on input amount, was done on the biotinylated dsDNA fraction bound to the beads using NEXTflex^TM^ kit amplification reagents. Amplified libraries were quantified by the Qubit dsDNA HS assay kit. Adaptor-dimers were removed by electrophoresis of the libraries on Novex 6% TBE gels (EC62655BOX, *Invitrogen*) at 120 V for 1 hour, and excising the smear above ∼150 bp. Gel slices containing libraries were shredded by centrifugation at 13,000 g for 3 minutes. Gel shreds were re-suspended in 500 μl crush and soak buffer (500 mM NaCl, 1.0 mM EDTA and 0.05% SDS) and incubated at 65°C for 2 hours on a thermomixer (1400 RPM for 15 s, rest for 45 s). Subsequently, the buffer was transferred into a Costar SpinX column (8161, *Corning Incorporated*) with two 1 cm glass pre-filters (1823010, *Whatman*). Columns were centrifuged at 13000 g for 1 min. DNA libraries in the flowthrough were precipitated at −20°C for 18 hours in ethanol with 0.3 M sodium acetate and 1 μl linear acrylamide (AM9520, *Ambion*). Purified libraries were further quantified and inspected on an Agilent 4200 TapeStation (*Agilent Technologies, Inc*). The libraries were sent for 100 bp SE sequencing by the Vincent J. Coates Genomics Sequencing Laboratory with a NovaSeq SP 100SR.

### Direct mRNA-seq

500 ng of poly(A)-selected mRNA was used as input for the Direct RNA Sequencing Kit (SQK-RNA002, *Oxford Nanopore Technologies*), used as directed with a modified reverse transcription (RT) step. Marathon reverse transcriptase (kindly gifted from Dr. Kathleen Collins) was used for the RT instead of Superscript III. The RT reaction was performed in 1X first strand buffer (20 mM Tris-HCl pH 7.5, 75 mM KCl, and 5 mM MgCl_2_), with 0.8 mM dNTPs, 8 mM DTT, and 20 μM Marathon reverse transcriptase. The RT reaction was incubated at 37°C for 50 min then 70°C for 10 min. Downstream steps were followed according to kit instructions. The library was loaded onto an R9.4.1 flow cell (FLO-MIN106, *Oxford Nanopore Technologies*) and sequenced on a minION (MIN-101B, *Oxford Nanopore Technologies*). MinKNOW (v22.05.5, *Oxford Nanopore Technologies*) was run without live base calling for 72 hours. Bases were called from fast5 files using Guppy (v6.0.1, *Oxford Nanopore Technologies*). Reads were aligned to the S288C reference genome (SacCer3) using the EPI2ME Desktop Agent (v3.5.6, *Oxford Nanopore Technologies*). Bam files were visualized directly in Integrated Genomics Viewer (IGV, *Broad Institute*).

### Chromatin Immunoprecipitation (ChIP)

Roughly 50 OD_600_ units of cells were collected in 50 ml conical tubes and fixed with 1% formaldehyde with periodic inversion for 15 min at room temperature. The crosslinking reaction was quenched with 125 mM glycine and tubes were gently agitated on a rocker platform (*Bellco Biotechnology*) at room temperature for 5 min. Cells were pelleted, washed with cold 1X PBS, then resuspended in 1 ml cold FA lysis buffer (50 mM Hepes pH 7.5, 150 mM NaCl, 1 mM EDTA, 1% Triton, 0.1% sodium deoxycholate) with 0.1% SDS and protease inhibitors (11836153001, *Roche*). In all cases, SDS was freshly added to buffers from a 20% stock solution. For ChIP-seq experiments, SK1 spike-in cells harboring a Snf2-3V5 allele (UB31748) that had been crosslinked in the same manner were added at a ratio of 1:5 prior to lysis. Roughly 500 μl Zirconia beads were added to cell suspension and cells were lysed by Beadbeater (Mini-Beadbeater-96, *Biospec Products*). Lysates were collected by centrifuging at 500 g for 1 minute at 4°C. Unbroken cells and debris were removed by microcentrifugation for 3 min at 2000 g. Supernatant was collected and centrifuged again for 20,000 g for 15 min resulting with pelleted chromatin. Pellets were resuspended in 1 ml FA lysis buffer with 0.1% SDS and protease inhibitors into falcon tubes containing 300 μl sonication beads. Samples were sonicated in a Bioruptor Pico (*Diagenode*) for 30 seconds on / 30 seconds off for 6 cycles to average fragment size of 150-400 bp. Samples were centrifuged once more at 20,000 g for 1 minute and the supernatant was carried forward to the immunoprecipitation (IP). From the isolated chromatin, 30 μl were set aside as input prior to IP.

For Snf2 immunoprecipitation, 25 μl of mouse anti-V5 agarose slurry (A7345, *MilliporeSigma*) were washed twice with 1 ml FA lysis buffer with 0.1% SDS. For each wash, the beads nutated at 4°C for 5 min and were subsequently pelleted by microcentrifugation (1000 g for 30 seconds). Sheared chromatin was added to the washed beads and the IP was incubated overnight with nutation at 4°C. IPs were washed at 4°C twice with FA lysis buffer and 0.1% SDS, twice with high salt buffer (FA lysis buffer with 0.1% SDS, and 250 mM NaCl) and twice with high detergent buffer (10 mM Tris pH 8, 250 mM LiCl, 0.5% NP-40, 0.5% sodium deoxycholate, and 1 mM EDTA). To IP and input samples, 130 μl TE (10 mM Tris pH 8, 1 mM EDTA) with 1% SDS was added. The precipitate was eluted from the beads by shaking at 450 RPM at 65°C in a thermomixer (*Eppendorf*) overnight.

Samples were cleaned up using QIAQuick PCR Purification Kit (28106, *QIAGEN*), diluted 1:10, and analyzed by qPCR with Absolute Blue qPCR Mix (AB4162B, *ThermoFisher Scientific*) using primer pairs directed against the *HNT1* locus or the *HMR* control locus. C_T_ mean values were first corrected for primer efficiency as calculated from standard curves performed on input samples for each primer pair, then corrected values corresponding to each *HNT1* primer pair were normalized over the corrected *HMR* signal. All ChIP experiments were performed on two biological replicates.

### Micrococcal nuclease digestion (MNase)

Roughly 50 OD_600_ units of cells were fixed in 1% formaldehyde with light shaking at RT for 15 minutes. Crosslinking was quenched by 125 mM of glycine for 5 minutes at RT. Cells were pelleted and washed with cold 1X PBS. For MNase-seq experiments, SK1 spike-in cells (UB31748) that had been crosslinked in the same manner were added at a ratio of 1:5 prior to spheroplasting. Cells were spheroplasted in 20 ml of Spheroplast Solution (1 M Sorbitol, 50 mM Tris pH 7.5, 10 mM β-mercaptoethanol) with 100 μL of 10 mg/ml zymolase until they appeared non-refractive and shadow-like after ∼12-15 minutes. Spheroplasted cells were resuspended in 2 ml MNase Digestion Buffer (1 M Sorbitol, 50 mM NaCl, 10 mM Tris pH 7.5, 5 mM MgCl_2_, 1 mM CaCl_2_, 0.075% NP-40, 0.5 mM spermidine, 1 mM β-mercaptoethanol). Digestions were performed with 600 μl of spheroplasts, 30 units of Exonuclease III (M0206S, *New England Biolabs*), and either 10, 20, or 40 units of MNase (LS004797, *Worthington*). 10 units of proteinase K (P8107S, *New England Biolabs*) were added to each digestion and undigested control. Crosslinks were then reversed by incubating samples at 65°C overnight and DNA was purified by 25:24:1 phenol:chloroform:isoamyl alcohol DNA extraction, followed by ethanol precipitation. Precipitated DNA was washed with 70% ethanol and resuspended in 1X NEBuffer #2 (B7002S, *New England Biolabs*) with 2 μl RNase A (20 mg/ml, 12091021, *Invitrogen*). Samples were incubated for 30 min and run on a 2% agarose gel to examine digestion efficiency. Of the samples digested with 10, 20, and 40 units of MNase, only the samples with a ratio of mononucleosomes to dinucleosomes closest to 80/20 were carried forward. Samples were once again purified by phenol/chloroform/isoamyl alcohol DNA extraction and ethanol precipitation, digests were eluted in 1X NEBuffer #3 (B7003S, *New England Biolabs*) and undigested gDNA was eluted in TE. Digests were then treated with 10 units alkaline phosphatase (CIP, P4978, *Sigma Aldrich*), incubated for 1 hour at 37°C, and size selected by running samples on a 1.8% LMT agarose gel at 120 V for 25 minutes at 4°C and gel extracting the mononucleosome band with a Monarch Gel Extraction Kit (T1020S, *New England Biolabs*). Purified samples were carried forward for qPCR or library preparation. All experiments were performed on two biological replicates.

### Sequencing libraries for DNA-, ChIP-, and MNase-Seq

DNA-seq, ChIP-seq, and MNase-seq libraries were generated with the NEXTflex^TM^ Rapid DNA-Seq Kit 2.0 Bundle (NOVA-5188-12, *Bioo Scientific*) according to manufacturer’s instructions. AMPure XP beads (A63881, *Beckman Coulter*) were used to select fragments between 200-800 bp (DNA-seq), 200-500 bp (ChIP-seq), or 150-250 bp (MNase-Seq). Libraries were quantified using the Agilent 4200 TapeStation (Agilent Technologies, Inc). Samples were submitted for 150 bp PE sequencing (DNA-seq) 100 bp SE sequencing (ChIP-seq), or 100 bp PE sequencing (MNase-seq) on a NovaSeq 6000 by the Vincent J. Coates Genomics Sequencing Laboratory.

### RNA extraction for RT-qPCR and RNA blotting

2 ml of cells were collected by centrifugation and snap frozen in liquid nitrogen. Cells were thawed on ice and resuspended in 400 μl TES buffer (10 mM Tris pH 7.5, 10 mM EDTA, 0.5% SDS). An equal volume of citrate-buffered acid phenol (pH 4.3, P4682, *Sigma-Aldrich*) was added to cells, and they were incubated at 65°C for 30 min in a Thermomixer (*Eppendorf*) shaking at 1400 RPM. After microcentrifugation (20,000 g for 10 min at 4°C) the aqueous phase was transferred to a second tube with 350 μl chloroform. The aqueous phase was again separated by microcentrifugation (20,000 g for 5 min at room temperature) and RNA was precipitated in 100% isopropanol with 350 mM sodium acetate (pH 5.2) overnight at –20°C. Pellets were washed with 80% ethanol and resuspended in DEPC water. Total RNA was quantified by Nanodrop.

### RNA blotting

10 μg of total RNA was denatured in a glyoxal/DMSO mix (1 M deionized glyoxal, 50% DMSO, 10 mM sodium phosphate (NaPi) buffer pH 6.5–6.8) at 70°C for 10 minutes. Denatured samples were mixed with loading buffer (10% glycerol, 2 mM NaPi buffer pH 6.5, 0.4% bromophenol blue) and separated on an agarose gel (1.1% agarose, 0.01 M NaPi buffer) for 3 hours at 100 V. RNA was transferred to a nitrocellulose membrane by capillary action using 10X SSC (1.5 M NaCl, 150 mM Na_3_Citrat-2H_2_O) overnight and crosslinked using a UV Crosslinker (*Stratagene*). rRNA bands were visualized using methylene blue staining. The membranes were blocked in ULTRAhyb Ultrasensitive Hybridization Buffer (AM8669, *ThermoFisher Scientific)* for 1 hour before overnight hybridization. Radioactive probes were synthesized using a MAXIScript T7 Transcription Kit (AM1314, *Invitrogen*). After synthesis, 1 μl of TURBO DNAse (2238G2, *Invitrogen*) was added and probes were incubated at 37°C for 10 min. Before spinning the probe mix through a hydrated NucAway column (AM10070, Invitrogen), 1 μl of EDTA (0.5M, pH 8) was added. Probes were eluted through the columns by spinning at 3,000 RPM for 3 min at room temperature, then added to hybridization tubes and incubated overnight. Membranes were washed twice in Low Stringency Buffer (2X SSC, 0.1% SDS) and three times in High Stringency Buffer (0.1X SSC, 0.1% SDS). All hybridization and wash steps were done at 68°C.

### Reverse transcription and qPCR

For each sample, 2.5 µg of RNA was treated with 1 unit of Turbo DNase using the TURBO DNA-free Kit (AM1907, *Invitrogen*) and incubated for 30 min at 37°C. 2.5 µl DNase inactivation reagent was added to stop the reaction and incubated for 5 min at room temperature. Samples were centrifuged for 2 minutes at 20,000 g and DNase-treated RNA was collected from the top layer.

2 μl of the treated RNA was then added to a 4.5 μl master mix containing 1 μl dNTPs (10 mM), 1 μl random hexamers (50 ng/μl), and nuclease-free water. Samples were incubated for 5 minutes at 65°C then cooled on ice for 1 minute.

cDNA was synthesized via reverse transcription using the SuperScript^TM^ III (18080044, *Invitrogen*) according to manufacturer’s instructions. Quantification was performed using 5.2 μl Absolute Blue qPCR Mix (AB4162B, *ThermoFisher Scientific*) and 4.8 μl of cDNA diluted 1:20 in nuclease-free water. Samples were run on a StepOnePlus (*Applied Biosystems*) qPCR machine in triplicate. The C_T_ mean from each primer pair directed against the experimental target was normalized using the ΔC_T_ method.

### Immunoblotting

To prepare lysates for SDS-PAGE, 2 OD_600_ units were pelleted at 2,000 g for 2 min at 4°C, resuspended in 1 ml of 5% trichloroacetic acid (SA433, *Thermo Fisher Scientific*) and incubated at 4°C for a minimum of 10 min. Samples were then spun at 2,000 g for 2 min at room temperature and cell pellets were washed by vortexing with 1 ml of 100% acetone. Samples were spun at maximum speed at room temperature for 5 min, acetone was pipetted off, and pellets were dried in a fume hood overnight. Once dried, pellets were resuspended in 100 μl of lysis buffer (50 mM Tris, pH 7.5, 1 mM EDTA, 3 mM DTT, and 1X cOmplete protease inhibitor cocktail [11836145001, *Roche*]) and lysed on a beadbeater for 5 min at room temperature with 100 μl of acid-washed glass beads. Next, 50 μl of 3X SDS buffer (18.75 mM Tris, pH 6.8, 6% β-mercaptoethanol, 30% glycerol, 9% SDS, 0.05% bromophenol blue) was added, and samples were boiled at 95°C for 5 min.

4–8 μl of samples and 3 μl of Precision Plus Protein Dual Color Standard (1610374, *Bio-Rad*) were loaded into 4–12% Bis-Tris Bolt gels (*Thermo Fisher Scientific*) and run at 150 V for 45 min. Protein was then transferred to 0.45 µm nitrocellulose membranes (*Bio-Rad*) with cold 1X Trans-Blot Turbo buffer in a semi-dry transfer apparatus (Trans-Blot Turbo Transfer System, *Bio-Rad*). Membranes were incubated at room temperature for 1 hour in PBS Odyssey Blocking Buffer (927-4000, *LI-COR*) and incubated in primary antibody solutions at 4°C overnight. Membranes were then washed three times in 1X PBS with 0.1% Tween-20 (PBS-T, 5 min per wash) at room temperature before incubating in secondary antibody solutions at room temperature for 2.5 hours. Membranes were washed three times in PBS-T at room temperature prior to imaging with the Odyssey system (*LI-COR Biosciences*).

All primary and secondary antibodies were diluted in PBS Odyssey Buffer with 0.1% Tween-20. Primary antibodies: mouse α-V5 antibody (R960-25, *Thermo Fisher Scientific*) used at a concentration of 1:2,000; rat α-tubulin (MCA78G, *Bio-Rad*) used at a concentration of 1:10,000. Secondary antibodies: goat α-mouse or α-rabbit secondary antibody conjugated to IRDye 800CW used at a concentration of 1:15,000 (926-32213, *LI-COR*); α-rabbit secondary conjugated to IRDye 680RD at a concentration of 1:15,000 (926-68071, *LI-COR*).

Immunoblot quantification was performed by quantifying signals from bands in Image Studio (*LI-COR*). For all blots quantified in this study, raw V5 signal was normalized to raw alpha-tubulin signal. For Hnt1 time course experiments, all normalized Hnt1-3V5 values from timepoints after t = 0 were then normalized to the levels at t = 0 to measure protein turnover.

#### Quantification and statistical analysis

Plotting and statistics for all bar charts, column plots, and scatterplots were performed using Prism Graphpad (v9.5.1).

### Identification of LUTI escape mutants

Fastq files were aligned to the SacCer3 reference genome with BWA-MEM (H. Li, 2013) and SAMBLASTER (v0.1.24, Faust and Hall, 2014) was used to remove discordant reads. SAMtools (v1.7, Danecek et al., 2021) was used to sort and index BAM files, then GATK (v4.1.7.0, Auwera and O’Connor, 2020) HaplotypeCaller was run on sorted BAM files with ‘-ERC GVCF’ to call variants. Genotyping was performed with GATK GenotypeGVCFs. Output VCF files were split into SNPs and indels with GATK SelectVariants. SNPs were filtered with GATK VariantFiltration “QD < 2.0” --filter-name “QD2” -filter “QUAL < 1000.0” --filter-name “QUAL1000” -filter “SOR > 3.0” --filter-name “SOR3” -filter “FS > 60.0” --filter-name “FS60” -filter “MQ < 40.0” --filter-name “MQ40” – filter “MQRankSum < -12.5” --filter-name “MQRankSum-12.5” -filter “ReadPosRankSum < -8.0” --filter-name “ReadPosRankSum-8” -filter “DP < 10” --filter-name “DP10”, while indels were filtered with GATK VariantFiltration “QD < 2.0” --filter-name “QD2” -filter “QUAL < 1000.0” --filter-name “QUAL1000” -filter “FS > 200.0” --filter-name “FS200” - filter “ReadPosRankSum < -20.0” --filter-name “ReadPosRankSum-20” -filter “DP < 10” --filter-name “DP10”. Next, a custom Python script was used to filter out reads in which the variant allele represents < 95% of total reads and output the filtered variants in BED format. Finally, variants that were also present in the parental control strain were filtered out using BEDtools subtract (v2.25.0, Quinlan and Hall, 2010). Mutations of interest were confirmed experimentally and tested for dominant/recessive phenotypes by backcrossing mutant strains to the parental control strain, phenotyping diploids and progeny, and performing Sanger-based sequencing to associate genotypes with phenotypes.

### mRNA-seq

Hisat2 (v2.1.0, Kim et al., 2019) was used to align reads to the sacCer3 reference genome (v64). Quantification of RNA as transcripts per million (TPM) was done with StringTie (v2.1.6, Pertea et al., 2015). Differential expression analysis was performed using DESeq2 (v1.36.0, Love et al., 2014) using default options. Hierarchical clustering was performed using Cluster3 (de Hoon et al., 2004) and visualized in Treeview (v1.2.0, Saldanha, 2004). TPM values for three biological replicates of each sample were log-adjusted and arrays were normalized in Cluster3 prior to clustering. Hierarchical clustering was performed on (1) genes based on Euclidian distance and (2) arrays using correlation-based similarity metrics. Average linkage was used as the hierarchical clustering method. Spearman heatmaps were produced in Jupyter Notebook (Kluyver et al., 2016) using seaborn (Waskom, 2021) and Matplotlib (Hunter, 2007). Average scores for Spearman’s rank sum correlation coefficient for each pairwise comparison are reported in the text.

### TL-seq and identification of TSS^DIST^/TSS^PROX^ targets

From the sequencing reads, the 3′ Illumina adaptor (AGATCGGAAGAGC) was trimmed using cutadapt with the–*minimum-length* option set to 20 bp (v2.3, Martin, 2011). From the 3′ trimmed output, the 5′ Illumina adaptor (CACTCTGAGCAATACC) was trimmed from reads with cutadapt. To select for reads representing the 5′ end of a transcript, only reads in which the 5′ adaptor was recognized and then trimmed were carried forward. Hisat2 was used to align reads to the sacCer3 reference genome. BAM files were loaded into CAGEr (v2.2.0, Haberle et al., 2015) and the CAGEexp object was created via getCTSS with correctSystematicG = FALSE and removeFirstG = FALSE. Next, reads were normalized by the normalizeTagCount function with method = c(“simpleTPM”). TSSs were then clustered with clusterCTSS with the following parameters: threshold = 1, thresholdIsTpm = TRUE, nrPassThreshold = 4, method = “distclu”, maxDist = 20, removeSingletons = TRUE, keepSingletonsAbove = 5. Consensus clusters were generated in a three-step process: (1) cumulativeCTSSdistribution was run on tagClusters with useMulticore = T, (2) quantilePositions was run on tagClusters with qLow = 0.1 and qUp = 0.9, and (3) aggregateTagClusters was run with tpmThreshold = 5, qLow = 0.1, qUp = 0.9, maxDist = 50. Consensus clusters were annotated using annotateConsensusClusters with an imported sacCer3 GFF file.

TPM tables from CAGEr-identified CTSS consensus clusters were exported as CSV files. Bed files used for metagene analysis centered on TSS positions were exported with rtracklayer and bedGraph files were exported with rtracklayer for visualization in IGV. Finally, Differential expression analysis was performed on consensus clusters using DESeq2 and results were exported for further analysis. For comparing wild-type samples in the untreated condition with DTT treatment, *SNF2* (n = 2) and *SWI3* (n = 2) wild-type control samples were combined for a total of n = 4 biological replicates per condition. This combination was determined based on genotypic and technical similarity between the wild-type samples (ρ ≥ 0.96 [untreated] and ≥ 0.98 [DTT]) to improve statistical power and simplify plotting. Lists of TSSs with significant upregulation (adjusted p-value < 0.05) in one or more LUTI escape mutant were curated in Excel and further manual curation was performed to identify sites with transcriptional readthrough. These sites were required to meet two criteria: (1) an upstream initiating transcript that extends through the TSS^PROX^ was visible in direct mRNA-seq reads and (2) the distal TSS was also called as a consensus cluster from CAGEr. Finally, sites that did not exhibit Snf2 binding as indicated by MACS2 peak calling in wild-type cells treated with DTT (see below) were eliminated from further analysis.

### ChIP-seq

Spike-in normalization was performed following the SNP-ChIP pipeline (Vale-Silva et al., 2019). First, reads from ChIP and input samples were aligned to a hybrid S288C-SK1 reference genome using Bowtie (v1.2.0, Langmead et al., 2009) with options -q -m 1 -v 0 -p 8 -S. Then, read counts mapping to each chromosome of the SK1 or S288C sequence were generated with SAMtools idxstats. Tables of read counts were used as input for spike-in normalization factor calculation using the R script published in (Vale-Silva et al., 2019). Supporting code and the S288C-SK1 hybrid reference genome is availble on Github (https://github.com/hochwagenlab/ChIPseq_functions).

Original fastq files were then re-aligned to the sacCer3 reference genome using Bowtie with options -q -p 8 -S. MACS2 (v2.1.1, Zhang et al., 2008) was used to call peaks with SPMR and normalize ChIP signals over input with bdgcmp. The output bedGraph files were then normalized by multiplying all coverage values with the spike-in normalization factor calculated as descibed above. Enrichment of Snf2 at DTT-induced alternative TSSs plotted in Figures 3B and S3A was calculated by averaging values from the normalized bedGraph files in the region between the TSS^DIST^ and TSS^PROX^ for each gene. For plotting enrichment, wild-type control strains *SNF2* and *SWI3* were combined as biological replicates on the bases of genotypic and technical similarity (ρ ≥ 0.98 [untreated] and ≥ 0.98 [DTT]). Normalized bedGraph files were converted to BigWig format for visualization in IGV and metagene analysis with deeptools (v3.5.1, Ramírez et al., 2016) bedGraphToBigWig. Metagene analysis was performed with deeptools computeMatrix reference-point using bed files corresponding to TSS regions generated from TL-seq analysis (see above) and visualized with plotProfile with --plotType heatmap.

### MNase-seq

Alignment and spike-in normalization of MNase reads were performed as described above for ChIP-seq normalization, using undigested gDNA from each sample as “input.” Aligned reads were filtered using deeptools alignmentSieve with options -- minFragmentLength 130 --maxFragmentLength 170. deeptools bamCoverage was used to generate bedGraph files, which were then multiplied by the spike-in normalization factor for each sample. Normalized bedGraph files were used to calculate nucleosome occupancy for plots in Figures S3D and S6C-D. Again, wild-type *SNF2* and *SWI3* samples were combined as wild-type biological replicates for plotting of nucleosome occupancy based on genotypic and technical similarity (ρ ≥ 0.96 [untreated] and ρ ≥ 0.93 [DTT]). Nucleosome fuzziness scores were calculated with DANPOS3 (K. Chen et al., 2013), which also outputs position coordinates. Determination of -1, +1, +2, and +5 nucleosome regions used for analysis in Figures 3I-J and S6D was performed as follows: BEDtools closest was run using a file containing TSS coordinates from TL-seq analysis (see above) and a file containing nucleosome coordinates from wild-type, unstressed cells output from DANPOS3. For the -1 nucleosome, options “-id -t first” were used. For the +1 nucleosome, options “-iu -t last” were used. For nucleosomes downstream of the +1, the previous nucleosome position was used as the reference coordinate file instead of the TSS coordinates (e.g. bedtools closest -a plus1.bed -b all_pos.bed -iu -t last > plus2.bed).

Plotting nucleosome occupancy in the NDR/NFR regions for Figures S3D and S6C was performed by analyzing sum occupancy, as calculated from spike-in normalized bedGraph files, in the region between the -1 and +1 nucleosomes. Averaged fuzziness and occupancy values are plotted for biological replicates (n = 4 for wild type and n = 2 for mutants). For nucleosome occupancy plotting, a cutoff score of 5 for normalized read depth in wild-type was applied. Individual fuzziness scores for the UPR TSS^DIST^/TSS^PROX^ targets are reported in Table S2. Normalized bedGraph files were converted to BigWig format for visualization in IGV and metagene analysis with deeptools bedGraphToBigWig. Metagene analyses were performed on BigWig files with deeptools computeMatrix reference-point using BED files corresponding to TSS regions generated from TL-seq analysis (see above) and visualized with plotProfile.

## Abbreviations

Swi/Snf: (Switch/Sucrose Nonfermenting)
LUTI: (Long Undecoded Transcript Isoform)
TF: (Transcription Factor)
NDR: (Nucleosome Depleted Region)
Pol II: (RNA Polymerase II)
UPR: (Unfolded Protein Response)
CDS: (Coding Sequence)
uORF: (Upstream Open Reading Frame)

## Resource availability

All reagents used in this study are available upon request from the corresponding author. Sequencing data generated in this study are available at NCBI GEO under the accession: GSE229952. The custom code used in filtering of DNA-seq data for LUTI escape mutants is available at: https://github.com/katemorse/LUTI_escape_filtering.git.

## Acknowledgements

We thank Gloria Brar, James Nuñez, Tina Sing, Amanda Su, Emily Powers, Grant King, Amy Tresenrider, Andrea Higdon, Ben Styler, Maia Reyes, Anthony Harris, Adriana Mendizabal, and Cyrus Ruediger for suggestions and comments on this manuscript. A special thanks to Molly Brothers, Pearl Omo-Sowho and Amy Tresenrider for their assistance with pilot LUTI escape selection strategies and Cal Milano from the Hochwagen lab for technical support with SNP-ChIP analysis. We also thank Kathleen Collins and her lab for kindly gifting us their MarathonRT reverse transcriptase. Finally, thanks to Michael Eisen, Gary Karpen, Arash Komeili, James Nuñez, Jeremy Thorner, and all members of the Brar and Ünal labs for valuable discussions. This work is supported by fund from the National Institutes of Health (F31 HD103399-01 to KM; R01 GM140005 to EÜ) and Astera Institute to EÜ.

## Author contributions

K.M., conceptualization, data curation, formal analysis, investigation, methodology, validation, visualization, and drafting of the manuscript. S.S., data curation, formal analysis, investigation, methodology, validation, visualization. E.Ü., conceptualization, methodology, visualization, supervision, project administration, funding acquisition, and drafting of the manuscript.

## REFERENCES

Arribere, J. A., & Gilbert, W. V. (2013). Roles for transcript leaders in translation and mRNA decay revealed by transcript leader sequencing. Genome Research, 23(6), 977–987. https://doi.org/10.1101/gr.150342.112

Auwera, G. van der, & O’Connor, B. D. (2020). Genomics in the cloud: Using Docker, GATK, and WDL in Terra (First edition). O’Reilly Media.

Bähler, J., Wu, J. Q., Longtine, M. S., Shah, N. G., McKenzie, A., Steever, A. B., Wach, A., Philippsen, P., & Pringle, J. R. (1998). Heterologous modules for efficient and versatile PCR-based gene targeting in Schizosaccharomyces pombe. Yeast (Chichester, England), 14(10), 943–951. https://doi.org/10.1002/(SICI)1097-0061(199807)14:10<943::AID-YEA292>3.0.CO;2-Y

Bai, L., & Morozov, A. V. (2010). Gene regulation by nucleosome positioning. Trends in Genetics, 26(11), 476–483. https://doi.org/10.1016/j.tig.2010.08.003

Bird, A. J., Gordon, M., Eide, D. J., & Winge, D. R. (2006). Repression of ADH1 and ADH3 during zinc deficiency by Zap1-induced intergenic RNA transcripts. The EMBO Journal, 25(24), 5726–5734. https://doi.org/10.1038/sj.emboj.7601453

Bowman, G. D. (2010). Mechanisms of ATP-dependent nucleosome sliding. Current Opinion in Structural Biology, 20(1), 73–81. https://doi.org/10.1016/j.sbi.2009.12.002

Brar, G. A., Yassour, M., Friedman, N., Regev, A., Ingolia, N. T., & Weissman, J. S. (2012). High-Resolution View of the Yeast Meiotic Program Revealed by Ribosome Profiling. Science, 335(6068), 552–557. https://doi.org/10.1126/science.1215110

Carrozza, M. J., Li, B., Florens, L., Suganuma, T., Swanson, S. K., Lee, K. K., Shia, W.– J., Anderson, S., Yates, J., Washburn, M. P., & Workman, J. L. (2005). Histone H3 Methylation by Set2 Directs Deacetylation of Coding Regions by Rpd3S to Suppress Spurious Intragenic Transcription. Cell, 123(4), 581–592. https://doi.org/10.1016/j.cell.2005.10.023

Cenik, B. K., & Shilatifard, A. (2021). COMPASS and SWI/SNF complexes in development and disease. Nature Reviews. Genetics, 22(1), 38–58. https://doi.org/10.1038/s41576-020-0278-0

Chen, J., Tresenrider, A., Chia, M., McSwiggen, D. T., Spedale, G., Jorgensen, V., Liao, H., Werven, F. J. van, & Ünal, E. (2017). Kinetochore inactivation by expression of a repressive mRNA. ELife, 6. https://doi.org/10.7554/eLife.27417

Chen, K., Xi, Y., Pan, X., Li, Z., Kaestner, K., Tyler, J., Dent, S., He, X., & Li, W. (2013). DANPOS: Dynamic analysis of nucleosome position and occupancy by sequencing. Genome Research, 23(2), 341–351. https://doi.org/10.1101/gr.142067.112

Chen, K., Yuan, J., Sia, Y., & Chen, Z. (2023). Mechanism of action of the SWI/SNF family complexes. Nucleus, 14(1), 2165604. https://doi.org/10.1080/19491034.2023.2165604

Cheng, Z., Otto, G. M., Powers, E. N., Keskin, A., Mertins, P., Carr, S. A., Jovanovic, M., & Brar, G. A. (2018). Pervasive, Coordinated Protein-Level Changes Driven by Transcript Isoform Switching during Meiosis. Cell, 172(5), 910–923.e16. https://doi.org/10.1016/j.cell.2018.01.035

Chia, M., Li, C., Marques, S., Pelechano, V., Luscombe, N. M., & van Werven, F. J. (2021). High-resolution analysis of cell-state transitions in yeast suggests widespread transcriptional tuning by alternative starts. Genome Biology, 22(1), 34. https://doi.org/10.1186/s13059-020-02245-3

Chia, M., Tresenrider, A., Chen, J., Spedale, G., Jorgensen, V., Ünal, E., & Werven, F. J. van. (2017). Transcription of a 5’ extended mRNA isoform directs dynamic chromatin changes and interference of a downstream promoter. ELife, 6. https://doi.org/10.7554/eLife.27420

Choi, J., Jeon, S., Choi, S., Park, K., & Seong, R. H. (2015). The SWI/SNF chromatin remodeling complex regulates germinal center formation by repressing Blimp-1 expression. Proceedings of the National Academy of Sciences of the United States of America, 112(7), E718–727. https://doi.org/10.1073/pnas.1418592112

Corey, L. L., Weirich, C. S., Benjamin, I. J., & Kingston, R. E. (2003). Localized recruitment of a chromatin-remodeling activity by an activator in vivo drives transcriptional elongation. Genes & Development, 17(11), 1392–1401. https://doi.org/10.1101/gad.1071803

Costa, P. J., & Arndt, K. M. (2000). Synthetic lethal interactions suggest a role for the Saccharomyces cerevisiae Rtf1 protein in transcription elongation. Genetics, 156(2), 535–547. https://doi.org/10.1093/genetics/156.2.535

Cox, J. S., & Walter, P. (1996). A novel mechanism for regulating activity of a transcription factor that controls the unfolded protein response. Cell, 87(3), 391–404. https://doi.org/10.1016/s0092-8674(00)81360-4

Danecek, P., Bonfield, J. K., Liddle, J., Marshall, J., Ohan, V., Pollard, M. O., Whitwham, A., Keane, T., McCarthy, S. A., Davies, R. M., & Li, H. (2021). Twelve years of SAMtools and BCFtools. GigaScience, 10(2), giab008. https://doi.org/10.1093/gigascience/giab008

Davie, J. K., & Kane, C. M. (2000). Genetic interactions between TFIIS and the Swi-Snf chromatin-remodeling complex. Molecular and Cellular Biology, 20(16), 5960–5973. https://doi.org/10.1128/MCB.20.16.5960-5973.2000

de Hoon, M. J. L., Imoto, S., Nolan, J., & Miyano, S. (2004). Open source clustering software. Bioinformatics, 20(9), 1453–1454. https://doi.org/10.1093/bioinformatics/bth078

Dutta, A., Gogol, M., Kim, J.-H., Smolle, M., Venkatesh, S., Gilmore, J., Florens, L., Washburn, M. P., & Workman, J. L. (2014). Swi/Snf dynamics on stress-responsive genes is governed by competitive bromodomain interactions. Genes & Development, 28(20), 2314–2330. https://doi.org/10.1101/gad.243584.114

Dutta, A., Sardiu, M., Gogol, M., Gilmore, J., Zhang, D., Florens, L., Abmayr, S. M., Washburn, M. P., & Workman, J. L. (2017). Composition and Function of Mutant Swi/Snf Complexes. Cell Reports, 18(9), 2124–2134. https://doi.org/10.1016/j.celrep.2017.01.058

Faust, G. G., & Hall, I. M. (2014). *SAMBLASTER*: Fast duplicate marking and structural variant read extraction. Bioinformatics, 30(17), 2503–2505. https://doi.org/10.1093/bioinformatics/btu314

Garg, A., Sanchez, A. M., Shuman, S., & Schwer, B. (2018). A long noncoding (lnc)RNA governs expression of the phosphate transporter Pho84 in fission yeast and has cascading effects on the flanking prt lncRNA and pho1 genes. The Journal of Biological Chemistry, 293(12), 4456–4467. https://doi.org/10.1074/jbc.RA117.001352

Gibson, D. G., Young, L., Chuang, R.-Y., Venter, J. C., Hutchison, C. A., & Smith, H. O. (2009). Enzymatic assembly of DNA molecules up to several hundred kilobases. Nature Methods, 6(5), 343–345. https://doi.org/10.1038/nmeth.1318

Gill, J. K., Maffioletti, A., García-Molinero, V., Stutz, F., & Soudet, J. (2020). Fine Chromatin-Driven Mechanism of Transcription Interference by Antisense Noncoding Transcription. Cell Reports, 31(5), 107612. https://doi.org/10.1016/j.celrep.2020.107612

Haberle, V., Forrest, A. R. R., Hayashizaki, Y., Carninci, P., & Lenhard, B. (2015). CAGEr: Precise TSS data retrieval and high-resolution promoterome mining for integrative analyses. Nucleic Acids Research, 43(8), e51–e51. https://doi.org/10.1093/nar/gkv054

Hainer, S. J., Pruneski, J. A., Mitchell, R. D., Monteverde, R. M., & Martens, J. A. (2011). Intergenic transcription causes repression by directing nucleosome assembly. Genes & Development, 25(1), 29–40. https://doi.org/10.1101/gad.1975011

Han, Y., Reyes, A. A., Malik, S., & He, Y. (2020). Cryo-EM structure of SWI/SNF complex bound to a nucleosome. Nature (London), 579(7799), 452–455. https://doi.org/10.1038/s41586-020-2087-1

Hangauer, M. J., Vaughn, I. W., & McManus, M. T. (2013). Pervasive transcription of the human genome produces thousands of previously unidentified long intergenic noncoding RNAs. PLoS Genetics, 9(6), e1003569. https://doi.org/10.1371/journal.pgen.1003569

Hartzog, G. A., Wada, T., Handa, H., & Winston, F. (1998). Evidence that Spt4, Spt5, and Spt6 control transcription elongation by RNA polymerase II in Saccharomyces cerevisiae. Genes & Development, 12(3), 357–369. https://doi.org/10.1101/gad.12.3.357

Hennig, B. P., Bendrin, K., Zhou, Y., & Fischer, T. (2012). Chd1 chromatin remodelers maintain nucleosome organization and repress cryptic transcription. EMBO Reports, 13(11), 997–1003. https://doi.org/10.1038/embor.2012.146

Hirschhorn, J. N., Brown, S. A., Clark, C. D., & Winston, F. (1992). Evidence that SNF2/SWI2 and SNF5 activate transcription in yeast by altering chromatin structure. Genes & Development, 6(12A), 2288–2298. https://doi.org/10.1101/gad.6.12a.2288

Hollerer, I., Barker, J. C., Jorgensen, V., Tresenrider, A., Dugast-Darzacq, C., Chan, L. Y., Darzacq, X., Tjian, R., Ünal, E., & Brar, G. A. (2019). Evidence for an Integrated Gene Repression Mechanism Based on mRNA Isoform Toggling in Human Cells. G3 (Bethesda, Md.), 9(4), 1045–1053. https://doi.org/10.1534/g3.118.200802

Hunter, J. D. (2007). Matplotlib: A 2D Graphics Environment. Computing in Science & Engineering, 9(3), 90–95. https://doi.org/10.1109/MCSE.2007.55

Jansen, A., & Verstrepen, K. J. (2011). Nucleosome Positioning in Saccharomyces cerevisiae. Microbiology and Molecular Biology Reviews, 75(2), 301–320. https://doi.org/10.1128/MMBR.00046-10

Jorgensen, V., Chen, J., Vander Wende, H., Harris, D. E., McCarthy, A., Breznak, S., Wong-Deyrup, S. W., Chen, Y., Rangan, P., Brar, G. A., Sawyer, E. M., Chan, L. Y., & Ünal, E. (2020). Tunable Transcriptional Interference at the Endogenous Alcohol Dehydrogenase Gene Locus in Drosophila melanogaster. G3 (Bethesda, Md.), 10(5), 1575–1583. https://doi.org/10.1534/g3.119.400937

Kadoch, C., & Crabtree, G. R. (2015). Mammalian SWI/SNF chromatin remodeling complexes and cancer: Mechanistic insights gained from human genomics. Science Advances, 1(5), e1500447. https://doi.org/10.1126/sciadv.1500447

Kim, D., Paggi, J. M., Park, C., Bennett, C., & Salzberg, S. L. (2019). Graph-based genome alignment and genotyping with HISAT2 and HISAT-genotype. Nature Biotechnology, 37(8), 907–915. https://doi.org/10.1038/s41587-019-0201-4

Kim, J. H., Lee, B. B., Oh, Y. M., Zhu, C., Steinmetz, L. M., Lee, Y., Kim, W. K., Lee, S. B., Buratowski, S., & Kim, T. (2016). Modulation of mRNA and lncRNA expression dynamics by the Set2-Rpd3S pathway. Nature Communications, 7, 13534. https://doi.org/10.1038/ncomms13534.

Kluyver, T., Ragan-Kelley, B., Pérez, F., Bussonnier, M., Frederic, J., Hamrick, J., Grout, J., Corlay, S., Ivanov, P., Abdalla, S., & Willing, C. (n.d.). Jupyter Notebooks—A publishing format for reproducible computational workflows.

Langmead, B., Trapnell, C., Pop, M., & Salzberg, S. L. (2009). Ultrafast and memory-efficient alignment of short DNA sequences to the human genome. Genome Biology, 10(3), R25. https://doi.org/10.1186/gb-2009-10-3-r25

Laprade, L., Winston, F., & Martens, J. A. (2004). Intergenic transcription is required to repress the Saccharomyces cerevisiae SER3 gene. Nature, 429(6991), 571–574. https://doi.org/10.1038/nature02538

Li, H. (n.d.). Aligning sequence reads, clone sequences and assembly contigs with BWA-MEM.

Li, M., Xia, X., Tian, Y., Jia, Q., Liu, X., Lu, Y., Li, M., Li, X., & Chen, Z. (2019). Mechanism of DNA translocation underlying chromatin remodelling by Snf2. Nature, 567(7748), 409–413. https://doi.org/10.1038/s41586-019-1029-2

Love, M. I., Huber, W., & Anders, S. (2014). Moderated estimation of fold change and dispersion for RNA-seq data with DESeq2. Genome Biology, 15(12), 550. https://doi.org/10.1186/s13059-014-0550-8

Lu, C., & Allis, C. D. (2017). SWI/SNF complex in cancer. Nature Genetics, 49(2), 178–179. https://doi.org/10.1038/ng.3779

Malagon, F., Tong, A. H., Shafer, B. K., & Strathern, J. N. (2004). Genetic interactions of DST1 in Saccharomyces cerevisiae suggest a role of TFIIS in the initiation-elongation transition. Genetics, 166(3), 1215–1227. https://doi.org/10.1534/genetics.166.3.1215

Martens, J. A., & Winston, F. (2002). Evidence that Swi/Snf directly represses transcription in S. cerevisiae. Genes & Development, 16(17), 2231–2236. https://doi.org/10.1101/gad.1009902

Martens, J. A., Wu, P.-Y. J., & Winston, F. (2005). Regulation of an intergenic transcript controls adjacent gene transcription in Saccharomyces cerevisiae. Genes & Development, 19(22), 2695–2704. https://doi.org/10.1101/gad.1367605

Martin, M. (2011). Cutadapt removes adapter sequences from high-throughput sequencing reads. EMBnet.Journal, 17(1), 10. https://doi.org/10.14806/ej.17.1.200

Menon, D. U., Shibata, Y., Mu, W., & Magnuson, T. (2019). Mammalian SWI/SNF collaborates with a polycomb-associated protein to regulate male germline transcription in the mouse. Development (Cambridge, England), 146(19), dev174094. https://doi.org/10.1242/dev.174094

Murphy, D. J., Hardy, S., & Engel, D. A. (1999). Human SWI-SNF Component BRG1 Represses Transcription of the c-fos Gene. Molecular and Cellular Biology, 19(4), 2724–2733.

Neigeborn, L., & Carlson, M. (1984). Genes affecting the regulation of SUC2 gene expression by glucose repression in Saccharomyces cerevisiae. Genetics, 108(4), 845–858. https://doi.org/10.1093/genetics/108.4.845

Olave, I. A., Reck-Peterson, S. L., & Crabtree, G. R. (2002). Nuclear actin and actin-related proteins in chromatin remodeling. Annual Review of Biochemistry, 71, 755– 781. https://doi.org/10.1146/annurev.biochem.71.110601.135507

Ottoz, D. S. M., Rudolf, F., & Stelling, J. (2014). Inducible, tightly regulated and growth condition-independent transcription factor in Saccharomyces cerevisiae. Nucleic Acids Research, 42(17), e130. https://doi.org/10.1093/nar/gku616

Peil, K., Värv, S., Ilves, I., Kristjuhan, K., Jürgens, H., & Kristjuhan, A. (2022). Transcriptional regulator Taf14 binds DNA and is required for the function of transcription factor TFIID in the absence of histone H2A.Z. The Journal of Biological Chemistry, 298(9), 102369. https://doi.org/10.1016/j.jbc.2022.102369

Pelechano, V., Wei, W., & Steinmetz, L. M. (2013). Extensive transcriptional heterogeneity revealed by isoform profiling. Nature, 497(7447), 127–131. https://doi.org/10.1038/nature12121

Pertea, M., Pertea, G. M., Antonescu, C. M., Chang, T.-C., Mendell, J. T., & Salzberg, S. L. (2015). StringTie enables improved reconstruction of a transcriptome from RNA-seq reads. Nature Biotechnology, 33(3), 290–295. https://doi.org/10.1038/nbt.3122

Pruneski, J. A., Hainer, S. J., Petrov, K. O., & Martens, J. A. (2011). The Paf1 Complex Represses SER3 Transcription in Saccharomyces cerevisiae by Facilitating Intergenic Transcription-Dependent Nucleosome Occupancy of the SER3 Promoter. Eukaryotic Cell, 10(10), 1283–1294. https://doi.org/10.1128/EC.05141-11

Quinlan, A. R., & Hall, I. M. (2010). BEDTools: A flexible suite of utilities for comparing genomic features. Bioinformatics, 26(6), 841–842. https://doi.org/10.1093/bioinformatics/btq033

Ramírez, F., Ryan, D. P., Grüning, B., Bhardwaj, V., Kilpert, F., Richter, A. S., Heyne, S., Dündar, F., & Manke, T. (2016). deepTools2: A next generation web server for deep-sequencing data analysis. Nucleic Acids Research, 44(W1), W160–W165. https://doi.org/10.1093/nar/gkw257

Rando, O. J., & Winston, F. (2012). Chromatin and Transcription in Yeast. Genetics, 190(2), 351–387. https://doi.org/10.1534/genetics.111.132266

Rawal, Y., Chereji, R. V., Qiu, H., Ananthakrishnan, S., Govind, C. K., Clark, D. J., & Hinnebusch, A. G. (2018). SWI/SNF and RSC cooperate to reposition and evict promoter nucleosomes at highly expressed genes in yeast. Genes & Development, 32(9–10), 695–710. https://doi.org/10.1101/gad.312850.118

Richmond, E., & Peterson, C. L. (1996). Functional Analysis of the DNA-Stimulated ATPase Domain of Yeast SWI2/SNF2. Nucleic Acids Research, 24(19), 3685– 3692. https://doi.org/10.1093/nar/24.19.3685

Rüegsegger, U., Leber, J. H., & Walter, P. (2001). Block of HAC1 mRNA translation by long-range base pairing is released by cytoplasmic splicing upon induction of the unfolded protein response. Cell, 107(1), 103–114. https://doi.org/10.1016/s0092-8674(01)00505-0

Sahu, R. K., Singh, S., & Tomar, R. S. (2021). The ATP-dependent SWI/SNF and RSC chromatin remodelers cooperatively induce unfolded protein response genes during endoplasmic reticulum stress. Biochimica et Biophysica Acta (BBA) – Gene Regulatory Mechanisms, 1864(11–12), 194748. https://doi.org/10.1016/j.bbagrm.2021.194748

Saldanha, A. J. (2004). Java Treeview—Extensible visualization of microarray data. Bioinformatics, 20(17), 3246–3248. https://doi.org/10.1093/bioinformatics/bth349

Schwabish, M. A., & Struhl, K. (2007). The Swi/Snf Complex Is Important for Histone Eviction during Transcriptional Activation and RNA Polymerase II Elongation In Vivo. Molecular and Cellular Biology, 27(20), 6987–6995. https://doi.org/10.1128/MCB.00717-07

Sehnal, D., Bittrich, S., Deshpande, M., Svobodová, R., Berka, K., Bazgier, V., Velankar, S., Burley, S. K., Koča, J., & Rose, A. S. (2021). Mol* Viewer: Modern web app for 3D visualization and analysis of large biomolecular structures. Nucleic Acids Research, 49(W1), W431–W437. https://doi.org/10.1093/nar/gkab314

Sen, P., Luo, J., Hada, A., Hailu, S. G., Dechassa, M. L., Persinger, J., Brahma, S., Paul, S., Ranish, J., & Bartholomew, B. (2017). Loss of Snf5 Induces Formation of an Aberrant SWI/SNF Complex. Cell Reports, 18(9), 2135–2147. https://doi.org/10.1016/j.celrep.2017.02.017

Shivaswamy, S., & Iyer, V. R. (2008). Stress-Dependent Dynamics of Global Chromatin Remodeling in Yeast: Dual Role for SWI/SNF in the Heat Shock Stress Response. Molecular and Cellular Biology, 28(7), 2221–2234. https://doi.org/10.1128/MCB.01659-07

Smith, C. L., & Peterson, C. L. (2005). A conserved Swi2/Snf2 ATPase motif couples ATP hydrolysis to chromatin remodeling. Molecular and Cellular Biology, 25(14), 5880–5892. https://doi.org/10.1128/MCB.25.14.5880-5892.2005

Smolle, M., Venkatesh, S., Gogol, M. M., Li, H., Zhang, Y., Florens, L., Washburn, M. P., & Workman, J. L. (2012). Chromatin remodelers Isw1 and Chd1 maintain chromatin structure during transcription by preventing histone exchange. Nature Structural & Molecular Biology, 19(9), 884–892. https://doi.org/10.1038/nsmb.2312

Su, A. J., Yendluri, S. C., & Ünal, E. (2023). *Control of meiotic entry by dual inhibition of a key mitotic transcription factor* [Preprint]. Genetics. https://doi.org/10.1101/2023.03.17.533246

Taggart, J., MacDiarmid, C. W., Haws, S., & Eide, D. J. (2017). Zap1-dependent transcription from an alternative upstream promoter controls translation of RTC4 mRNA in zinc-deficient Saccharomyces cerevisiae. Molecular Microbiology, 106(5), 678–689. https://doi.org/10.1111/mmi.13851

Tatip, S., Taggart, J., Wang, Y., MacDiarmid, C. W., & Eide, D. J. (2020). Changes in transcription start sites of Zap1-regulated genes during zinc deficiency: Implications for HNT1 gene regulation. Molecular Microbiology, 113(1), 285–296. https://doi.org/10.1111/mmi.14416

Tresenrider, A., Chia, M., van Werven, F. J., & Ünal, E. (2022). Long undecoded transcript isoform (LUTI) detection in meiotic budding yeast by direct RNA and transcript leader sequencing. STAR Protocols, 3(1), 101145. https://doi.org/10.1016/j.xpro.2022.101145

Tresenrider, A., Morse, K., Jorgensen, V., Chia, M., Liao, H., van Werven, F. J., & Ünal, E. (2021). Integrated genomic analysis reveals key features of long undecoded transcript isoform-based gene repression. Molecular Cell, 81(10), 2231–2245.e11. https://doi.org/10.1016/j.molcel.2021.03.013

Turegun, B., Baker, R. W., Leschziner, A. E., & Dominguez, R. (2018). Actin-related proteins regulate the RSC chromatin remodeler by weakening intramolecular interactions of the Sth1 ATPase. Communications Biology, 1, 1. https://doi.org/10.1038/s42003-017-0002-6

Vale-Silva, L. A., Markowitz, T. E., & Hochwagen, A. (2019). SNP-ChIP: a versatile and tag-free method to quantify changes in protein binding across the genome. In BMC Genomics (Vol. 20, Issue 1). https://doi.org/10.1186/s12864-018-5368-4

Van Dalfsen, K. M., Hodapp, S., Keskin, A., Otto, G. M., Berdan, C. A., Higdon, A., Cheunkarndee, T., Nomura, D. K., Jovanovic, M., & Brar, G. A. (2018). Global Proteome Remodeling during ER Stress Involves Hac1-Driven Expression of Long Undecoded Transcript Isoforms. Developmental Cell, 46(2), 219–235.e8. https://doi.org/10.1016/j.devcel.2018.06.016

Vander Wende, H. M., Gopi, M., Onyundo, M., Medrano, C., Adanlawo, T., & Brar, G. A. (2023). Meiotic resetting of the cellular Sod1 pool is driven by protein aggregation, degradation, and transient LUTI-mediated repression. The Journal of Cell Biology, 222(3), e202206058. https://doi.org/10.1083/jcb.202206058

Venkatesh, S., & Workman, J. L. (2015). Histone exchange, chromatin structure and the regulation of transcription. Nature Reviews. Molecular Cell Biology, 16(3), 178–189. https://doi.org/10.1038/nrm3941

Wang, X., Hou, J., Quedenau, C., & Chen, W. (2016). Pervasive isoform-specific translational regulation via alternative transcription start sites in mammals. Molecular Systems Biology, 12(7), 875. https://doi.org/10.15252/msb.20166941

Waskom, M. (2021). seaborn: Statistical data visualization. Journal of Open Source Software, 6(60), 3021. https://doi.org/10.21105/joss.03021

Werven, F. J. van Neuert, G., Hendrick, N., Lardenois, A., Buratowski, S., Oudenaarden, A. van Primig, M., & Amon, A. (2012). Transcription of Two Long Noncoding RNAs Mediates Mating-Type Control of Gametogenesis in Budding Yeast. Cell, 150(6), 1170–1181. https://doi.org/10.1016/j.cell.2012.06.049

Yang, X., Zaurin, R., Beato, M., & Peterson, C. L. (2007). Swi3p controls SWI/SNF assembly and ATP-dependent H2A-H2B displacement. Nature Structural & Molecular Biology, 14(6), 540–547. https://doi.org/10.1038/nsmb1238

Yen, K., Vinayachandran, V., Batta, K., Koerber, R. T., & Pugh, B. F. (2012). Genome-wide Nucleosome Specificity and Directionality of Chromatin Remodelers. Cell, 149(7), 1461–1473. https://doi.org/10.1016/j.cell.2012.04.036

Zhang, Y., Liu, T., Meyer, C. A., Eeckhoute, J., Johnson, D. S., Bernstein, B. E., Nusbaum, C., Myers, R. M., Brown, M., Li, W., & Liu, X. S. (2008). Model-based Analysis of ChIP-Seq (MACS). Genome Biology, 9(9), R137. https://doi.org/10.1186/gb-2008-9-9-r137

Zhu, Y., Rowley, M. J., Böhmdorfer, G., & Wierzbicki, A. T. (2013). A SWI/SNF Chromatin-Remodeling Complex Acts in Noncoding RNA-Mediated Transcriptional Silencing. Molecular Cell, 49(2), 298–309. https://doi.org/10.1016/j.molcel.2012.11.011

