## Supplemental Figure and Table Legends for "Swi/Snf Chromatin Remodeling Regulates Transcriptional Interference and Gene Repression"

### SUPPLEMENTAL FIGURE LEGENDS

**Figure S1. Candidate-based approach for LUTI escape phenotypes.**

**A.** *HIS3*$^{LUTI}$ serial dilution and spotting growth assay. Cells were plated on synthetic complete media lacking histidine with 200 µM 3-AT (left) or 200 µM 3-AT and 25 nM β-estradiol (right) and grown for 72 h at 30°C before imaging. Strains (from top to bottom): UB17314, UB13670, UB33427, UB33424, UB33482, and UB34040. **B.** *ADE2*$^{LUTI}$ color assay. Cells were streaked onto a YPD plate lacking supplemental adenine and grown for 24 h at 30°C before imaging. Strains: *ade2-1* (UB7), *ADE2*$^{LUTI}$ (UB21565), *swp82Δ ADE2*$^{LUTI}$ (UB30456), *snf11Δ ADE2*$^{LUTI}$ (UB30507), *swi3Δ ADE2*$^{LUTI}$ (UB27673), and *snf2Δ ADE2*$^{LUTI}$ (UB27669). **(C)** Growth curves for cells grown in rich media with 2% dextrose (YPD, left) or 2% sucrose (right). Absorbance readings at 600 nm collected every 15 minutes for 24 h is plotted for the following strains: WT (UB4784), *swp82Δ* (UB31746), *snf11Δ* (UB31807), *snf2Δ* (UB20060), and *swi3Δ* (UB27896). **(D)** *HIS3*$^{LUTI}$ serial dilution and spotting growth assay. Cells were plated on synthetic complete media lacking histidine with 200 µM 3-AT (left) or 200 µM 3-AT and 25 nM β-estradiol (right) and grown for 72 h at 30°C before imaging. Strains (from top to bottom): UB24051, UB22912, UB24057, UB23996, UB24000, UB29782, UB29784, UB23999, and UB30085. **(E)** *ADE2*$^{LUTI}$ color assay. Cells were streaked onto a YPD plate lacking supplemental adenine and grown for 24 h at 30°C before imaging. Strains, all harboring *ADE2*$^{LUTI}$: control (UB22912), *isw1Δ* (UB24057), *isw2Δ* (UB23996), *chd1Δ* (UB24000), *isw1Δ isw2Δ* (UB29782), *isw1Δ chd1Δ* (UB29784), *isw2Δ chd1Δ* (UB23999), and *isw1Δ isw2Δ chd1Δ* (UB30085)

**Figure S2. Swi/Snf loss-of-function assays and Spearman correlation heatmaps.**

**A.** Growth curves for cells grown in rich media with 2% dextrose (YPD). Absorbance readings at 600 nm collected every 15 minutes for 24 h is plotted for the following strains: *SWI3* (UB19205, left, black), *swi3-E815X* (UB19209, left, blue), *swi3Δ* (UB27896, left, gray), *SNF2* (UB28914, right, black), *snf2-Q935R* (UB28922, right, purple), *snf2-Q928K* (UB28915, right, orange), and *snf2Δ* (UB29781, right, gray). **B.** RT-qPCR measuring relative abundance of *SRG1* (left) and *SER3* (right) mRNA in Swi/Snf mutants compared to wild-type cells (n = 3). Student's t test was performed on each mutant-to-wild type comparison (two-tailed, p = 0.3490 [*swi3-E815X*], p = 0.1718 [*snf2-W935R*], p = 0.0050 [snf2-*Q928K*], p = 0.1331 [*swi3Δ*], p = 0.0035 [*snf2Δ*]). Strains are the same as listed in (A). **C.** Same as (B), but for *SER3*. Student's t test was performed on each mutant-to-wild type comparison (two-tailed, p = 0.4956 [*swi3-E815X*], p = 0.0871 [*snf2-W935R*], p < 0.0001 [snf2-*Q928K*], p = 0.0018 [*swi3Δ*], p = 0.0058 [*snf2Δ*]). **D.** Heatmaps were generated using Spearman's rank correlation coefficient values from mRNA sequencing data portrayed in Figure 2D. For relevant pairwise comparisons (WT vs. null, null vs. LUTI escape mutant, and LUTI escape mutant vs. WT) the Spearman's rank sum correlation coefficient is reported in the main text.

**Figure S3. Single-gene validation and analysis of mutant ChIP-seq and MNase-seq signals.**

**A.** Snf2 ChIP-seq signals plotted for the 12 genes identified by the strategy outlined in Figure 3A. The average Snf2 binding levels are compared between wild type (n = 4) and LUTI escape mutants (n = 2) in the untreated condition (paired t test, two-tailed, p = 0.2263 [*swi3-E815X*], p = 0.1839 [*snf2-W935R*], p = 0.0385 [*snf2-Q928K*]) and DTT-treated conditions (p = 0.0398 [*swi3-E815X*], p = 0.0514 [*snf2-W935R*], p = 0.0300 [*snf2-Q928K*]). Strains: wild type (UB30387 and UB30070), *swi3-E815X* (UB30071), *snf2-W935R* (UB30391), and *snf2-Q928K* (UB30389). **B.** Genome browser snapshot. TL-Seq and direct mRNA-seq data from this study are overlaid with ribosome footprints from Van Dalfsen *et. al* 2018 for the genes *ADI1* and *ODC2*. For *ODC2*, the distal isoform is expressed in the unstressed and DTT condition, but expressed increases with DTT treatment. For *ADI1,* the distal isoform is expressed exclusively in response to DTT treatment. In both cases, there is uORF translation within the 5′ leader sequence of the distal isoform. **C.** Quantification of immunoblot portrayed in Figure 3G for two biological replicates. Student's t test (two-tailed) revealed a significant increase in Odc2 protein levels upon deletion of the $ODC2^{DIST}$ promoter in both untreated and DTT-treated cells (p =0.0036 [untreated], p = 0.0023 [DTT]), whereas Adi1 protein levels were significantly increased with deletion of the $ADI1^{DIST}$ promoter in DTT-treated condition only (p = 0.7281 [untreated], p = 0.0016 [DTT]. Erg27 protein levels were not found to be significantly different with deletion of the $ERG27^{DIST}$ promoter in either condition (p = 0.1988 [untreated], p = 0.3291 [DTT]). **D.** MNase signals, normalized with spike-in correction (see Materials and Methods for details), for the $TSS^{PROX}$ NDR/NFR among the 12 genes identified by the strategy in Figure 3A are plotted for wild type cells (n = 4) and LUTI escape mutants (n = 2) that were untreated or treated with 5 mM DTT. Strains are the same as in (A). A paired t test was performed on wild-type nucleosome occupancy comparing untreated to DTT-treated cells (two-tailed, p = 0.0445) and each mutant-to-wild type comparison in untreated conditions (p = 0.9629 [*swi3-E815X*], p = 0.4263 [*snf2-W935R*], p = 0.5048 [*snf2-Q928K*]) or with 5 mM DTT treatment (p = 0.1441 [*swi3-E815X*], p = 0.2004 [*snf2-W935R*], p = 0.2293 [*snf2-Q928K*]).

**Figure S4. *HNT1* and *HAC1* RNA blots.**

**A.** RNA blot probed for the *HNT1* CDS in wild type cells (UB32339) or cells lacking the *HNT1*$^{LUTI}$ promoter (*LUTIΔ*, UB32342). Cells were collected at 0, 15, 30, and 60 minutes after treatment with 5 mM DTT for RNA extraction. rRNA bands were detected by methylene blue staining. **B.** RNA blot probed for the *HAC1* CDS in cells that were untreated or treated with DTT for 1 h. Strains: *SNF2* (UB30152), *snf2-W935R* (UB30156), *snf2-Q928K* (UB30154), *SWI3* (UB24251), and *swi3-E815X* (UB24253).

**Figure S6. Gene expression and chromatin analysis for Swi/Snf targets and control genes.**

**A.** Scatterplot with TPM values, plotted on logarithmic scale, depicting expression of Swi/Snf target genes (n = 250) Left: *swi3-E815X* mutant (UB19209) compared to *SWI3* control cells (UB19205). Middle: *snf2-W35R* mutant (UB28922) compared to *SNF2* control cells (UB28914). Right: *snf2-Q928K* mutant (UB28915) compared to *SNF2* control cells (UB28914). One of three biological replicates is shown. **B.** Same as (A), but for non-Swi/Snf regulated control genes (n = 250). **C.** Quantification of nucleosome occupancy within the NDR for non-regulated control genes and Swi/Snf targets. $Log_2$(fold change) in occupancy was calculated from MNase-seq read depth after spike-in normalization for *swi3-E815X* (n = 2, UB30071), *snf2-W935R* (n = 2, UB30391), and *snf2-Q928K* (n = 2, UB30389) mutants relative to wildtype (n = 4, UB30070 and UB30387). Occupancy differences between control genes and Swi/Snf target genes is analyzed by Mann-Whitney U test (two tailed, p = 0.0018 [*swi3-E815X*], p = 0.0081 [*snf2-W935R*], p < 0.0001 [*snf2-Q928K*]. **D.** Same as (C), but for the +1, +2, and + 5 nucleosomes. Differences are analyzed by Mann-Whitney U test (two tailed, p = 0.2054 [*swi3-E815X*, +1], p = 0.9986 [*snf2-W935R*, +1], p = 0.6092 [*snf2-Q928K*, +1], p = 0.0425 [*swi3-E815X, +2*], p = 0.0006 [*snf2-W935R*, +2], p = 0.0001 [*snf2-Q928K*, +2], p = 0.1654 [*swi3-E815X*, +5], p = 0.0112 [*snf2-W935R*, +5], p = 0.7697 [*snf2-Q928K*, +5]).

#### SUPPLEMENTAL TABLE LEGENDS

**Table S1.**

Genotypes and phenotypes of mutants uncovered from LUTI escape selection strategy to identify regulators of LUTI-based interference.

† Growth was scored on media lacking histidine with 25 nM β-estradiol and 200 μM 3-amino-1,2,4-triazole (3-AT). The parent control strain used for genetic selection fails to grow in these conditions.

†† Color was scored on yeast extract peptone with 2% dextrose (YPD) media with limiting adenine. The parent control strain used for screening is red in these conditions.

††† Conservation determined by whether the mutation affects a conserved residue (in the case of missense mutations) or region (in the case of nonsense or frameshift mutations) of the human protein homolog.

**Table S2.**

TL-Seq and MNase-seq quantification for the 12 genes analyzed in Figure 3. Data reported for the untreated and DTT-treated conditions include: TPM values from wild-type cells for the distal and proximal isoforms, $\log_2$ (fold change) expression and adjusted p-value for each mutant-to-wild type comparison or wild type untreated-to-DTT output from DESeq2, qualitative observation of uORF translation within the distal isoform from Van Dalfsen et al., 2018 data, and nucleosome fuzziness scores output from DANPOS3.

**Table S3.**

Lists of Swi/Snf targets and control non-target genes analyzed in Figure 6. List of genes eliminated from the Swi/Snf target list based on their differential expression in *swi3-E815X* and *snf2-W935R* mutants. Normalization factors calculated from SNP-ChIP and SNP-MNase spike-in normalization for ChIP-seq and MNase-seq experiments.

**Table S4.**

List of yeast strains and genotypes used in this study.

**Table S5.**

List of oligonucleotides used in this study for quantitative PCR, RNA blotting, and TL-seq library preparation.

**Table S6.**

List of plasmids generated in this study.
