## Supplemental Figures for "Swi/Snf Chromatin Remodeling Regulates Transcriptional Interference and Gene Repression"

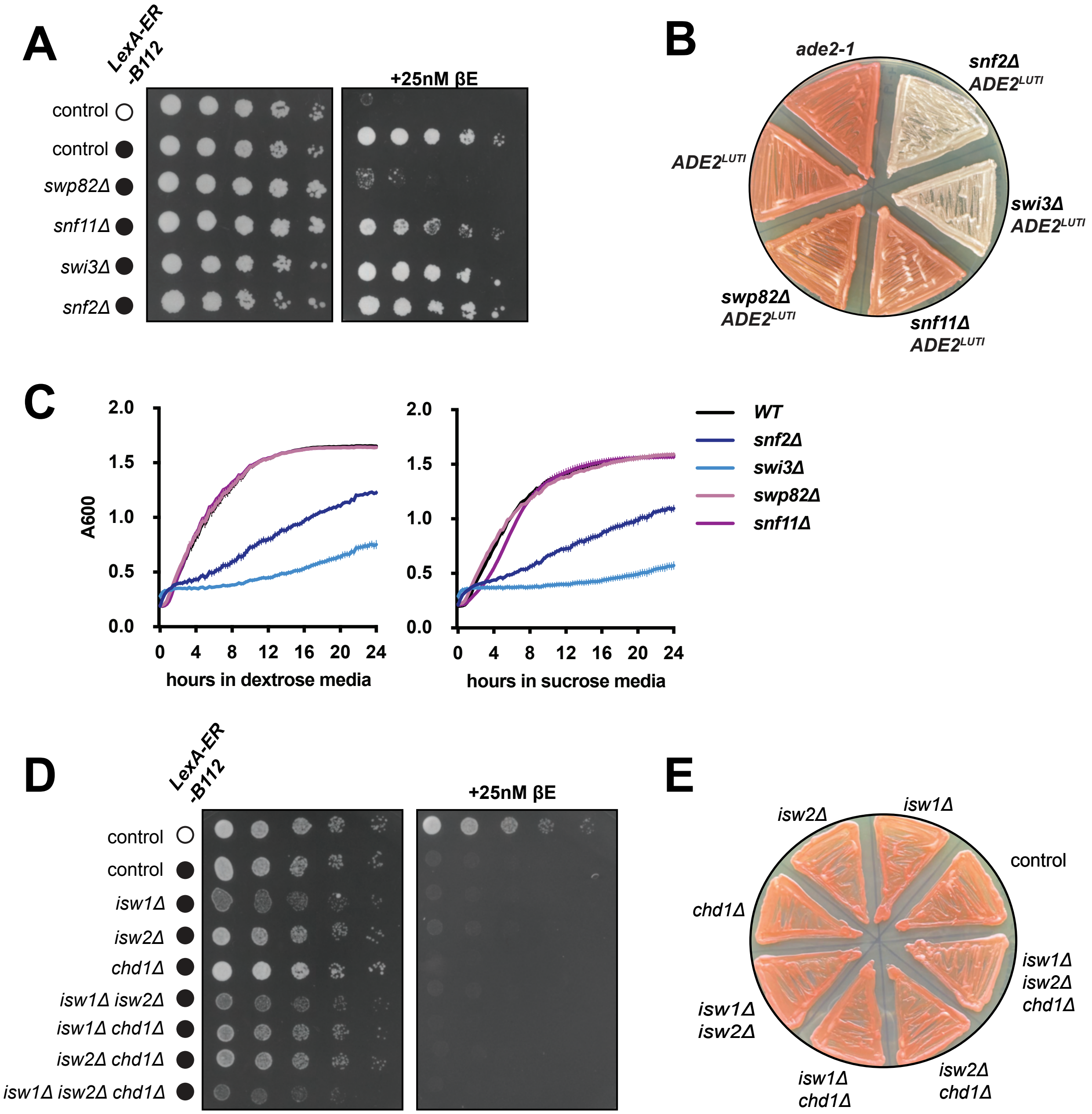


Morse et al_Figure S1

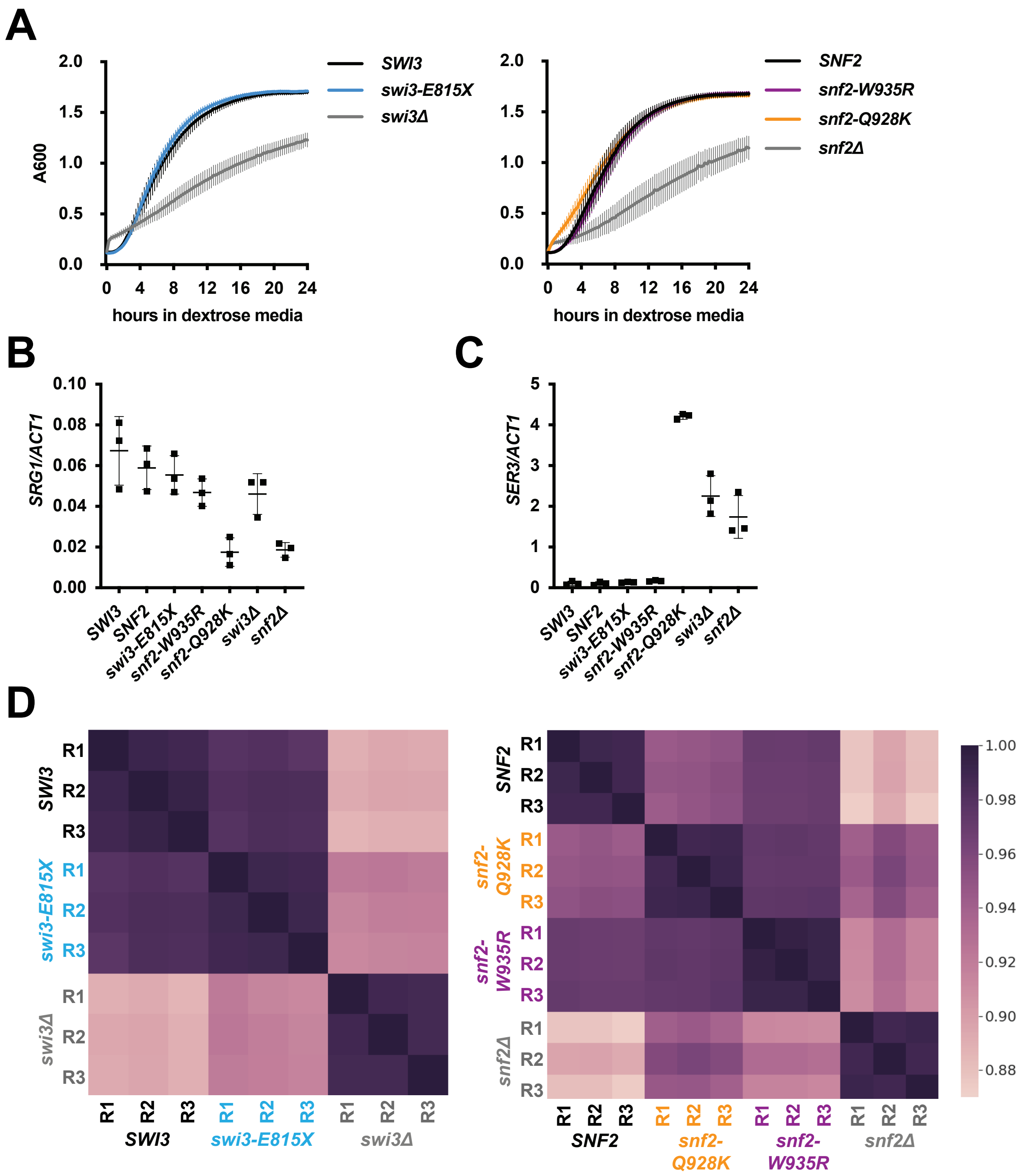


Morse et al_Figure S2

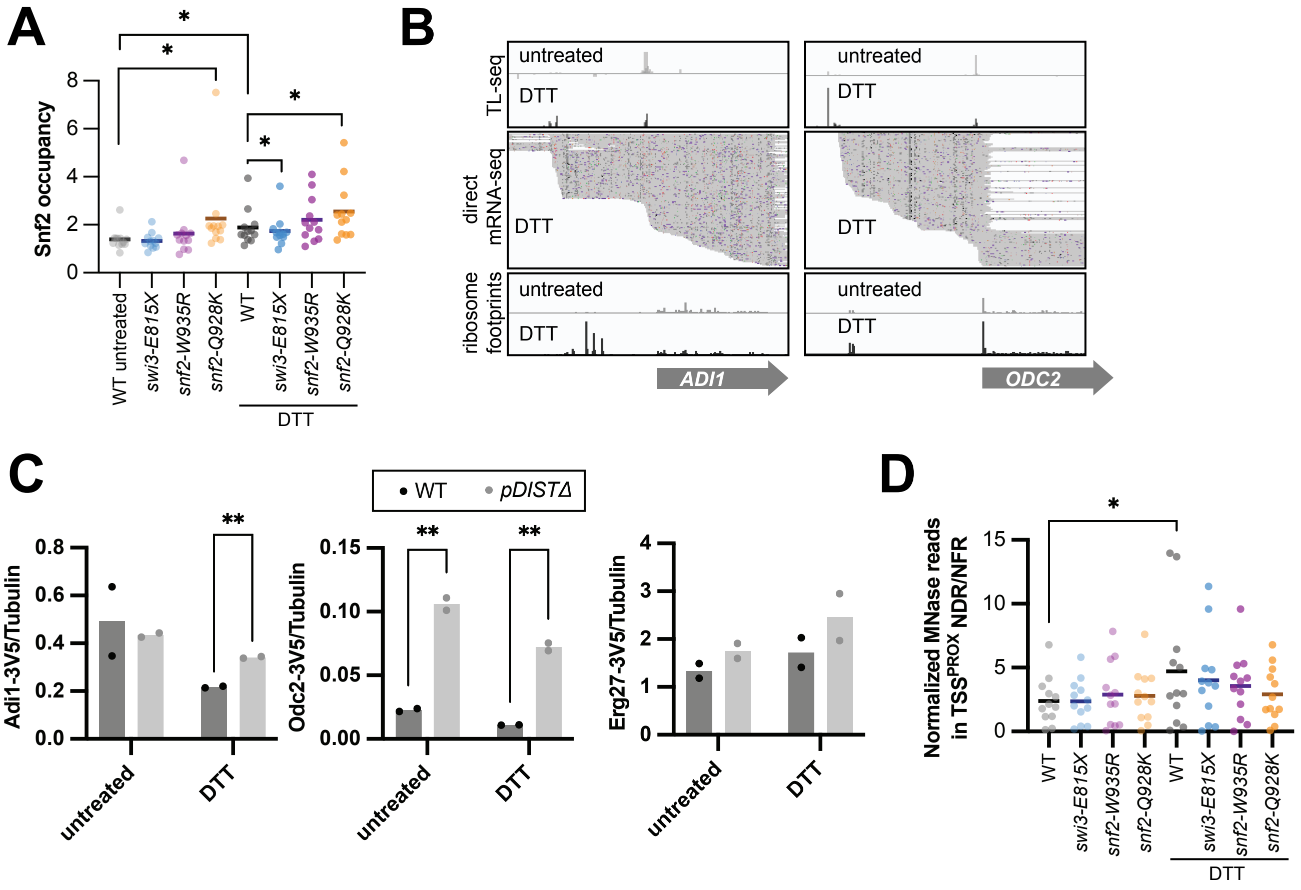


Morse et al_Figure S3

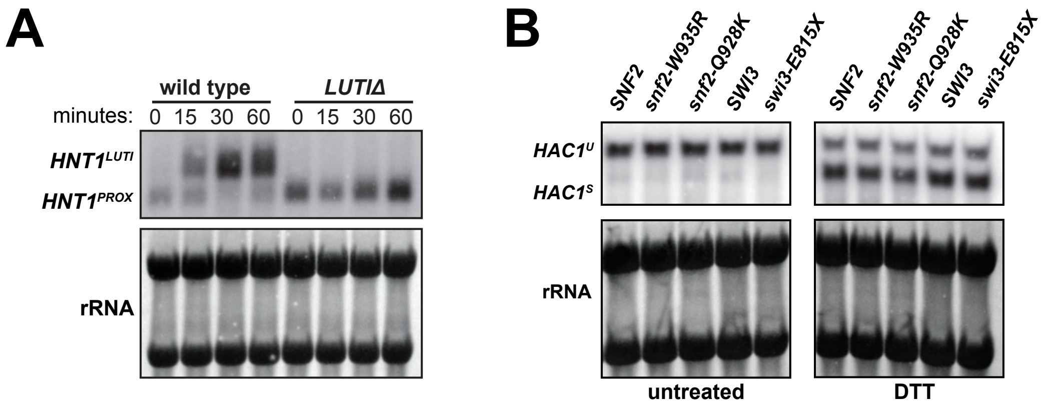


### Morse et al_Figure S4

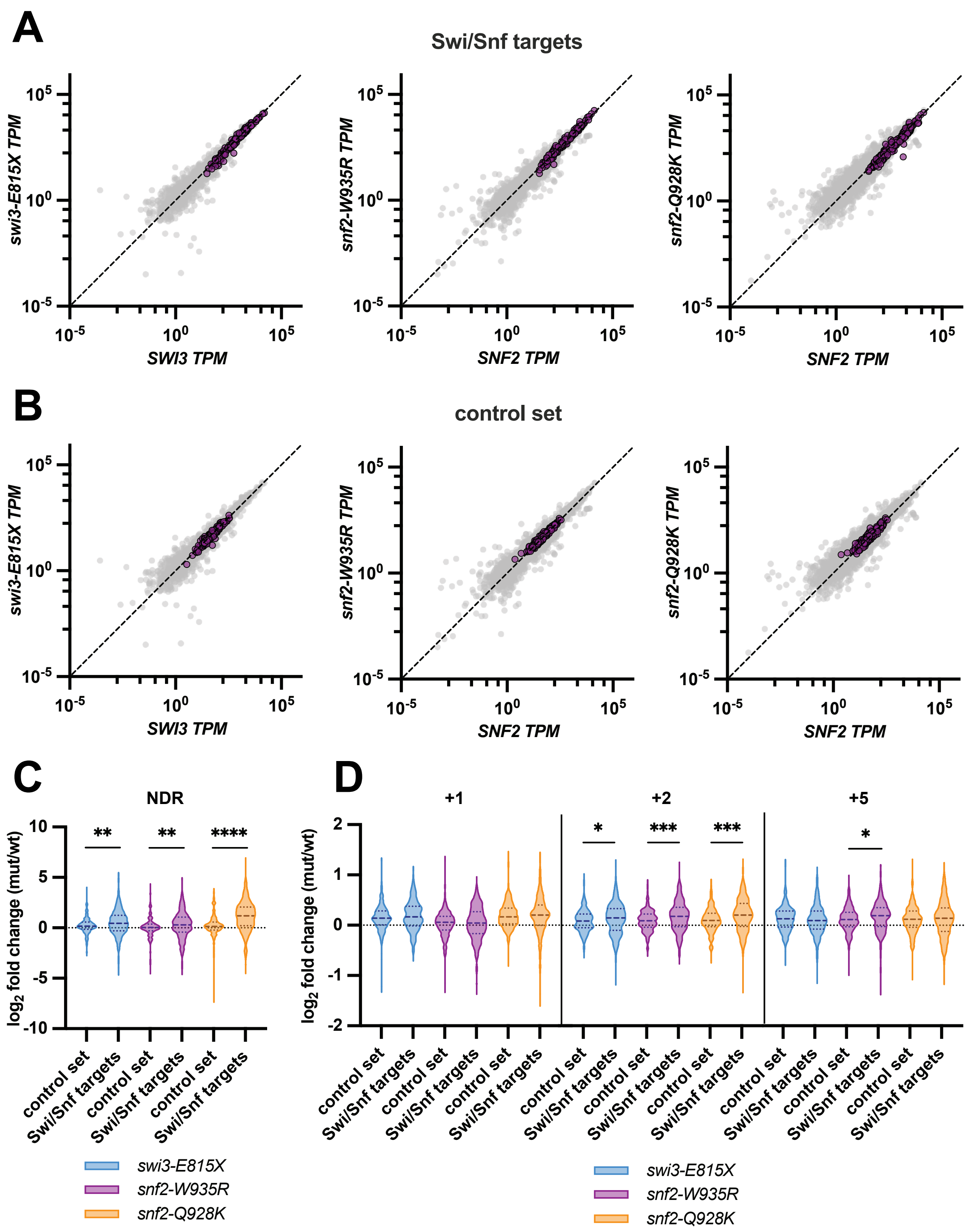


**Morse et al_Figure S6**
