## Supplementary material for "Swi/Snf Chromatin Remodeling Regulates Transcriptional Interference and Gene Repression": Table S1

| Gene | Mutation | Disruption of *HIS3^LUTI^*-based repression  (growth) ^†^ | Disruption of *ADE2^LUTI^*^-^based repression  (color) ^††^ | LUTI escape Phenotype | Conserved?^†††^ |
| --- | --- | --- | --- | --- | --- |
| *SNF2* | Frameshift at His 412 | ++ | pink | recessive | yes |
| *SNF2* | Frameshift at Lys 651 | + | cream | recessive | yes |
| *SNF2* | Gln 928 > Lys | ++ | cream | dominant negative | yes |
| *SNF2* | Trp 935 > Arg | ++ | pink | recessive | yes |
| *SNF2* | Glu 973 > Stop | + | cream | recessive | yes |
| *SNF5* | Gln 225 > Stop | ++ | pink | recessive | yes |
| *SNF5* | Gln 267 > Stop | + | pink | recessive | yes |
| *SNF12* | Gln 226 > Stop | + | cream | recessive | yes |
| *SWI1* | Ser 764 > Stop | ++ | pink | recessive | no |
| *SWI3* | Glu 815 > Stop | ++ | pink | recessive | no |
| *SNF6* | Arg 135 > Stop | + | cream | recessive | no |

**Table S1. List of LUTI escape mutants.**
