## Supplementary material for "Swi/Snf Chromatin Remodeling Regulates Transcriptional Interference and Gene Repression": Table S4

**Supplemental Table 5. Genotypes for the strains used in this study**

| **Strain** | **Genotype** |
| --- | --- |
| UB7, *W303* wild type | *MATa, ade2-1, leu2-3, ura3, trp1-1, his3-11,15, can1-100, GAL, psi+* |
| UB4784,  ADE+ *W303* wild type | *MATa, ADE2, leu2-3, ura3, trp1-1, his3-11,15, can1-100, GAL, phi+* |
| UB13670 | *MATa, ADE2, leu2-3, ura3, trp1-1, his3-11,15, can1-100, GAL, psi+ his3::3xLexO-NDC80leader::HIS3* |
| UB17314 | *MATa, ADE2, leu2-3, ura3, trp1-1, his3-11,15, can1-100, GAL, psi+ trp1::pGPD1-LexA-ER-HA-B112::TRP1 his3::3xLexO-NDC80leader::HIS3* |
| UB19205 | *MATa, ADE2, leu2-3, ura3, trp1-1, his3-11,15, can1-100, GAL, phi+ swi3::KANMX leu2::SWI3::LEU2* |
| UB19209 | *MATa, ADE2, leu2-3, ura3, trp1-1, his3-11,15, can1-100, GAL, phi+ swi3::KANMX leu2::swi3-E815X::LEU2* |
| UB20060 | *MATa, ADE2, leu2-3, ura3, trp1-1, his3-11,15, can1-100, GAL, phi+ snf2::KANMX* |
| UB21565 | *MATa, ade2-1, leu2-3, ura3, trp1-1, his3-11,15, can1-100, GAL, psi+*  *ura3::pTEF1-ADE2luti::URA3* |
| UB22912 | *MATa, ade2-1, leu2-3, ura3, trp1-1, his3-11,15, can1-100, GAL, psi+ trp1::pGPD1-LexA-ER-HA-B112::TRP1 ura3::pTEF1-ADE2luti::URA3 his3::3xLexO-NDC80leader::HIS3* |
| UB23545 | *MATa, ade2-1, leu2-3, ura3, trp1-1, his3-11,15, can1-100, GAL, psi+ swi3::KANMX leu2::swi3-E815X::LEU2 ura3::pTEF1-ADE2luti::URA3* |
| UB23996 | *MATa, ade2-1, leu2-3, ura3, trp1-1, his3-11,15, can1-100, GAL, psi+ trp1::pGPD1-LexA-ER-HA-B112::TRP1 ura3::pTEF1-ADE2luti::URA3 his3::3xLexO-NDC80leader::HIS3 isw2::HygB* |
| UB23999 | *MATa, ade2-1, leu2-3, ura3, trp1-1, his3-11,15, can1-100, GAL, psi+ trp1::pGPD1-LexA-ER-HA-B112::TRP1 ura3::pTEF1-ADE2luti::URA3 his3::3xLexO-NDC80leader::HIS3 chd1::HygB isw2::HygB* |
| UB24000 | *MATa, ade2-1, leu2-3, ura3, trp1-1, his3-11,15, can1-100, GAL, psi+ trp1::pGPD1-LexA-ER-HA-B112::TRP1 ura3::pTEF1-ADE2luti::URA3 his3::3xLexO-NDC80leader::HIS3 chd1::HygB* |
| UB24051 | *MATa, ade2-1, leu2-3, ura3, trp1-1, his3-11,15, can1-100, GAL, psi+ ura3::pTEF1-ADE2luti::URA3 his3::3xLexO-NDC80leader::HIS3* |
| UB24057 | *MATa, ade2-1, leu2-3, ura3, trp1-1, his3-11,15, can1-100, GAL, psi+ trp1::pGPD1-LexA-ER-HA-B112::TRP1 ura3::pTEF1-ADE2luti::URA3 his3::3xLexO-NDC80leader::HIS3 isw1::HygB* |
| UB24251 | *MATa, ADE2, leu2-3, ura3, trp1-1, his3-11,15, can1-100, GAL, phi+ hnt1::HNT1-3V5::KANMX swi3::KANMX leu2::SWI3::LEU2* |
| UB24253 | *MATa, ADE2, leu2-3, ura3, trp1-1, his3-11,15, can1-100, GAL, phi+ hnt1::HNT1-3V5::KANMX swi3::KANMX leu2::swi3-E815X::LEU2* |
| UB24301 | *MATa, ADE2, leu2-3, ura3, trp1-1, his3-11,15, can1-100, GAL, phi+  his3::3xLexO-NDC80leader::HIS3 trp1::pGPD1-LexA-ER-HA-B112::TRP1 swi3::KANMX leu2::swi3-E815X::LEU2* |
| UB27669 | *MATa, ade2-1, leu2-3, ura3, trp1-1, his3-11,15, can1-100, GAL, psi+ ura3::pTEF1-ADE2luti::URA3*  *snf2::KanMX* |
| UB27673 | *MATa, ade2-1, leu2-3, ura3, trp1-1, his3-11,15, can1-100, GAL, psi+ ura3::pTEF1-ADE2luti::URA3*  *swi3::KanMX* |
| UB27896 | *MATa, ADE2, LEU2, ura3, trp1-1, his3-11,15, can1-100, GAL, phi+ swi3::KANMX* |
| UB28096 | *MATa, ADE2, leu2-3, ura3, trp1-1, his3-11,15, can1-100, GAL, phi+ swi3::KANMX leu2::swi3-E815X::LEU2 dst1::HygB* |
| UB28907 | *MATa, ADE2, leu2-3, ura3, trp1-1, his3-11,15, can1-100, GAL, phi+ SNF2::LEU2 trp1::pGPD1-LexA-ER-HA-B112::TRP1 his3::3xLexO-NDC80leader::HIS3* |
| UB28911 | *MATa, ADE2, leu2-3, ura3, trp1-1, his3-11,15, can1-100, GAL, phi+ snf2::KANMX SNF2::LEU2 trp1::pGPD1-LexA-ER-HA-B112::TRP1 his3::3xLexO-NDC80leader::HIS3* |
| UB28914 | *MATa, ADE2, leu2-3, ura3, trp1-1, his3-11,15, can1-100, GAL, phi+ snf2::KANMX SNF2::LEU2* |
| UB28915 | *MATa, ADE2, leu2-3, ura3, trp1-1, his3-11,15, can1-100, GAL, phi+ snf2::KANMX snf2-Q928K::LEU2* |
| UB28919 | *MATa, ADE2, leu2-3, ura3, trp1-1, his3-11,15, can1-100, GAL, phi+ snf2::KANMX snf2-Q928K::LEU2 trp1::pGPD1-LexA-ER-HA-B112::TRP1 his3::3xLexO-NDC80leader::HIS3* |
| UB28922 | *MATa, ADE2, leu2-3, ura3, trp1-1, his3-11,15, can1-100, GAL, phi+ snf2::KANMX snf2-W935R::LEU2* |
| UB28923 | *MATa, ade2-1, leu2-3, ura3, trp1-1, his3-11,15, can1-100, GAL, phi+ snf2::KANMX snf2-W935R::LEU2 ura3::pTEF1-ADE2luti::URA3* |
| UB28925 | *MATa, ADE2, leu2-3, ura3, trp1-1, his3-11,15, can1-100, GAL, phi+ snf2::KANMX snf2-W935R::LEU2 trp1::pGPD1-LexA-ER-HA-B112::TRP1 his3::3xLexO-NDC80leader::HIS3* |
| UB29161 | *MATa, ADE2, leu2-3, ura3, trp1-1, his3-11,15, can1-100, GAL, phi+  SNF2-VL-3V5::HISMX* |
| UB29166 | *MATa, ADE2, leu2-3, ura3, trp1-1, his3-11,15, can1-100, GAL, phi+ snf2-W935R::LEU2 trp1::pGPD1-LexA-ER-HA-B112::TRP1 his3::3xLexO-NDC80leader::HIS3* |
| UB29170 | *MATa, ADE2, leu2-3, ura3, trp1-1, his3-11,15, can1-100, GAL, phi+ snf2-Q928K::LEU2 trp1::pGPD1-LexA-ER-HA-B112::TRP1 his3::3xLexO-NDC80leader::HIS3* |
| UB29188 | *MATa, ADE2, LEU2, ura3, trp1-1, his3-11,15, can1-100, GAL, phi+  trp1::pGPD1-LexA-ER-HA-B112::TRP1 his3::3xLexO-NDC80leader::HIS3* |
| UB29385 | *MATa, ADE2, LEU2, ura3, TRP1, his3-11,15, can1-100, GAL, phi+ his3::3xLexO-NDC80leader::HIS3* |
| UB29694 | *MATa, ADE2, leu2-3, ura3, trp1-1, his3-11,15, can1-100, GAL, phi+  his3::3xLexO-NDC80leader::HIS3 trp1::pGPD1-LexA-ER-HA-B112::TRP1 leu2::swi3-E815X::LEU2* |
| UB29781 | *MATa, ADE2, LEU2, ura3, trp1-1, his3-11,15, can1-100, GAL, phi+ snf2::KANMX* |
| UB29782 | *MATa, ade2-1, leu2-3, ura3, trp1-1, his3-11,15, can1-100, GAL, psi+ trp1::pGPD1-LexA-ER-HA-B112::TRP1 ura3::pTEF1-ADE2luti::URA3 his3::3xLexO-NDC80leader::HIS3 isw1::HygB isw2::HygB* |
| UB29784 | *MATa, ade2-1, leu2-3, ura3, trp1-1, his3-11,15, can1-100, GAL, psi+ trp1::pGPD1-LexA-ER-HA-B112::TRP1 ura3::pTEF1-ADE2luti::URA3 his3::3xLexO-NDC80leader::HIS3 isw1::HygB chd1::HygB* |
| UB29791 | *MATa, ADE2, leu2-3, ura3, trp1-1, his3-11,15, can1-100, GAL, phi+ swi3::KANMX leu2::SWI3::LEU2 his3::3xLexO-NDC80leader::HIS3 trp1::pGPD1-LexA-ER-HA-B112::TRP1* |
| UB29792 | *MATa, ADE2, leu2-3, ura3, trp1-1, his3-11,15, can1-100, GAL, phi+ leu2::SWI3::LEU2 his3::3xLexO-NDC80leader::HIS3 trp1::pGPD1-LexA-ER-HA-B112::TRP1* |
| UB30034 | *MATa, ade2-1, leu2-3, ura3, trp1-1, his3-11,15, can1-100, GAL, phi+ ura3::pTEF1-ADE2luti::URA3 snf2::KANMX SNF2::LEU2* |
| UB30070 | *MATa, ADE2, leu2-3, ura3, trp1-1, his3-11,15, can1-100, GAL, phi+  SNF2-VL-3V5::HISMX swi3::KANMX leu2::SWI3::LEU2* |
| UB30071 | *MATa, ADE2, leu2-3, ura3, trp1-1, his3-11,15, can1-100, GAL, phi+  SNF2-VL-3V5::HISMX swi3::KANMX leu2::swi3-E815X::LEU2* |
| UB30085 | *MATa, ade2-1, leu2-3, ura3, trp1-1, his3-11,15, can1-100, GAL, psi+ trp1::pGPD1-LexA-ER-HA-B112::TRP1 ura3::pTEF1-ADE2luti::URA3 his3::3xLexO-NDC80leader::HIS3 isw1::HygB chd1::HygB isw2::HygB* |
| UB30152 | *MATa, ADE2, leu2-3, ura3, trp1-1, his3-11,15, can1-100, GAL, psi+ snf2::KANMX SNF2::LEU2 hnt1::HNT1-3V5::KANMX* |
| UB30154 | *MATa, ADE2, leu2-3, ura3, trp1-1, his3-11,15, can1-100, GAL, psi+ snf2::KANMX snf2-Q928K::LEU2 hnt1::HNT1-3V5::KANMX* |
| UB30156 | *MATa, ADE2, leu2-3, ura3, trp1-1, his3-11,15, can1-100, GAL, phi+ snf2::KANMX snf2-W935R::LEU2 hnt1::HNT1-3V5::KANMX* |
| UB30185 | *MATa, ade2-1, leu2-3, ura3, trp1-1, his3-11,15, can1-100, GAL, phi+ snf2::KANMX snf2-Q928K::LEU2 ura3::pTEF1-ADE2luti::URA3* |
| UB30190 | *MATa, ade2-1, leu2-3, ura3, trp1-1, his3-11,15, can1-100, GAL, psi+ swi3::KANMX leu2::SWI3::LEU2 ura3::pTEF1-ADE2luti::URA3* |
| UB30387 | *MATa, ADE2, leu2-3, ura3, trp1-1, his3-11,15, can1-100, GAL, phi+ snf2::KANMX leu2::LEU2::SNF2-3V5::HISMX* |
| UB30389 | *MATa, ADE2, leu2-3, ura3, trp1-1, his3-11,15, can1-100, GAL, phi+ snf2::KANMX leu2::LEU2::snf2-Q928K-3V5::HISMX* |
| UB30391 | *MATa, ADE2, leu2-3, ura3, trp1-1, his3-11,15, can1-100, GAL, phi+ snf2::KANMX leu2::LEU2::snf2-W935R-3V5::HISMX* |
| UB30456 | *MATa, ade2-1, leu2-3, ura3, trp1-1, his3-11,15, can1-100, GAL, psi+ trp1::pGPD1-LexA-ER-HA-B112::TRP1 ura3::pTEF1-ADE2luti::URA3 his3::3xLexO-NDC80leader::HIS3*  *swp82::KanMX* |
| UB30507 | *MATa, ade2-1, leu2-3, ura3, trp1-1, his3-11,15, can1-100, GAL, psi+ trp1::pGPD1-LexA-ER-HA-B112::TRP1 ura3::pTEF1-ADE2luti::URA3 his3::3xLexO-NDC80leader::HIS3*  *snf11::KanMX* |
| UB31746 | *MATalpha, ADE2, leu2-3, ura3, trp1-1, his3-11,15, can1-100, GAL, psi+ swp82::KanMX* |
| UB31748 | *MATa, ho::LYS2 lys2 ura3 leu2::hisG his3::hisG trp1::hisG*  *Snf2-3V5::HISMX*  SK1 |
| UB31807 | *MATa, ADE2, leu2-3, ura3, trp1-1, his3-11,15, can1-100, GAL, phi+ snf11::KanMX* |
| UB32339 | *MATa, ADE2, leu2-3, ura3, trp1-1, his3-11,15, can1-100, GAL, phi+ trp1:: HNT1-3V5::TRP1 HNT1::KANMX* transgene contains HNT1luti TSS and UPRE sites  Endogenous locus lacks HNT1 5’ regulatory region and CDS |
| UB32342 | *MATa, ADE2, leu2-3, ura3, trp1-1, his3-11,15, can1-100, GAL, phi+ trp1:: HNT1(lutiΔ)-3V5::TRP1 HNT1::KANMX* transgene lacks HNT1luti TSS and UPRE sites  Endogenous locus lacks HNT1 5’ regulatory region and CDS |
| UB33424 | *MATa, ADE2, leu2-3, ura3, trp1-1, his3-11,15, can1-100, GAL, psi+ trp1::pGPD1-LexA-ER-HA-B112::TRP1 his3::3xLexO-NDC80leader::HIS3*  *snf11:KANMX* |
| UB33427 | *MATa, ADE2, leu2-3, ura3, trp1-1, his3-11,15, can1-100, GAL, psi+ trp1::pGPD1-LexA-ER-HA-B112::TRP1 his3::3xLexO-NDC80leader::HIS3*  *swp82::KANMX* |
| UB33482 | *MATa, ADE2, leu2-3, ura3, trp1-1, his3-11,15, can1-100, GAL, psi+ trp1::pGPD1-LexA-ER-HA-B112::TRP1 his3::3xLexO-NDC80leader::HIS3*  *swi3::KANMX* |
| UB34040 | *MATa, ADE2, leu2-3, ura3, trp1-1, his3-11,15, can1-100, GAL, psi+ trp1::pGPD1-LexA-ER-HA-B112::TRP1 his3::3xLexO-NDC80leader::HIS3*  *snf2::KANMX* |
| UB36048 | *MATa, ADE2, leu2-3, ura3, trp1-1, his3-11,15, can1-100, GAL, phi+ trp1::HNT1(pLUTI-CYC1t)-3V5::TRP1 HNT1::KANMX* CYC1 terminator inserted between luti and prox TSS  Endogenous locus lacks HNT1 5’ regulatory region and CDS |
| UB36182 | *MATa, ADE2, leu2-3, ura3, trp1-1, his3-11,15, can1-100, GAL, phi+ dst1::HygB swi3::KANMX leu2::SWI3::LEU2* |
| UB36185 | *MATa, ADE2, leu2-3, ura3, trp1-1, his3-11,15, can1-100, GAL, phi+ dst1::HygB snf2::KANMX SNF2::LEU2* |
| UB36186 | *MATa, ADE2, leu2-3, ura3, trp1-1, his3-11,15, can1-100, GAL, phi+ dst1::HygB snf2::KANMX snf2-W935R::LEU2* |
| UB36188 | *MATa, ADE2, leu2-3, ura3, trp1-1, his3-11,15, can1-100, GAL, phi+ dst1::HygB snf2::KANMX snf2-Q928K::LEU2* |
| UB36511 | *MATa, ADE2, leu2-3, ura3, trp1-1, his3-11,15, can1-100, GAL, phi+ trp1::ADI1-3V5::TRP1* |
| UB36513 | *MATa, ADE2, leu2-3, ura3, trp1-1, his3-11,15, can1-100, GAL, phi+ trp1::ADI1(pDISTΔ)-3V5::TRP1* |
| UB36515 | *MATa, ADE2, leu2-3, ura3, trp1-1, his3-11,15, can1-100, GAL, phi+ trp1::ODC2-3V5::TRP1* |
| UB36521 | *MATa, ADE2, leu2-3, ura3, trp1-1, his3-11,15, can1-100, GAL, phi+ trp1::ODC2(pDISTΔ)-3V5::TRP1* |
| UB36594 | *MATa, ADE2, leu2-3, ura3, trp1-1, his3-11,15, can1-100, GAL, phi+ trp1::ERG27-3V5::TRP1* |
| UB36596 | *MATa, ADE2, leu2-3, ura3, trp1-1, his3-11,15, can1-100, GAL, phi+ trp1::ERG27(pDISTΔ)-3V5::TRP1* |
