## Supplementary material for "Swi/Snf Chromatin Remodeling Regulates Transcriptional Interference and Gene Repression": Table S5

**Supplemental Table 6. Primers used for quantitative PCR, TL-seq, and RNA blotting**

| **Primer Name** | **Sequence from 5′ to 3′** |
| --- | --- |
| SRG1_RTqPCR_F | GGTTTTCTGAGCGGGATGAA |
| SRG1_RTqPCR_R | CCTTATCCTCTGCTCCCTCC |
| SER3_RTqPCR_F | ATCTGCCCCACAATTTGCTG |
| SER3_RTqPCR_R | GCTTTCACGGCTTGGATCAA |
| ACT1_RTqPCR_F | GTACCACCATGTTCCCAGGTATT |
| ACT1_RTqPCR_R | AGATGGACCACTTTCGTCGT |
| 5oligocap (TL-seq 5’ adapter) | dCdAdCdTdCdTrGrArGrCrArArUrArCrC |
| Second strand biotinylated oligo (TL-seq) | GCAC/iBiodT/GCACTCTGAGCAATACC |
| V5_probe_F | CTAGTGGATCCAGGTAAACCTAT |
| V5_probe_R | taatacgactcactataggCCAGTCCTAATAGAGGATTAGG |
| HNT1_probe_F | CATGGTGCGAAGTTGCATG |
| HNT1_probe_R | taatacgactcactataggCCACCCTACAATCAAACCAC |
| HNT1_RTqPCR_F  (also used in ChIP and MNase qPCR assays, anneals between LUTI and PROX TSS) | TGGTGCGAATCGTTACAGAA |
| HNT1_RTqPCR_R  (also used in ChIP and MNase qPCR assays, anneals between LUTI and PROX TSS) | AATGCTTCAGTAGGGCGGTA |
| HNT1_LUTIpromoter1_F | GCAAGGACCCAAATAGGAG |
| HNT1_LUTIpromoter1_R | GATTTACCGGTGTTTCCTTTG |
| HNT1_LUTIpromoter2_F | CAAAGGAAACACCGGTAAATC |
| HNT1_LUTIpromoter2_R | CTGTAGACAAGTGTCAATTCAACC |
| HNT1_LUTI_TSS_F | GGTTGAATTGACACTTGTCTACAG |
| HNT1_LUTI_TSS_R | TTCTGTAACGATTCGCACCA |
| HNT1_PROX_TSS_F | TACCGCCCTACTGAAGCATT |
| HNT1_PROX_TSS_R | CGTGCTGATTGTCCTTTTACTTC |
| HNT1_ORF1_F | GAAGTAAAAGGACAATCAGCACG |
| HNT1_ORF1_R | CAAGCGTAGCAGGAGCAGAC |
| HNT1_ORF2_F | GTCTGCTCCTGCTACGCTTG |
| HNT1_ORF2_R | CATCATTGGTGTGAGTGTAAGC |
| HNT1_ORF3_F | GCTTACACTCACACCAATGATG |
| HNT1_ORF3_R | ATGGAATTTCGCCTGTTGCATAG |
| HMR_F | ACGATCCCCGTCCAAGTTATG |
| HMR_R | CTTCAAAGGAGTCTTAATTTCCCTG |
| PHO5_F | CCATTTGGGATAAGGGTAAACATC |
| PHO5_R | AGAGATGAAGCCATACTAACCTCG |
| HAC1_probe_F | GCAGTCAGGTTTGAATTCATTTGAATTGAATGATTTC |
| HAC1_probe_R | taatacgactcactataggGCCTCTTCTTCTTCGGTTGAAGTAGCACACAC |
| SOD1_probe_F | CAACCACTGTCTCTTACGAGATCGC |
| SOD1_probe_R | taatacgactcactataggCACCATTTTCGTCCGTCTTTACG |
