## Supplementary material for "Swi/Snf Chromatin Remodeling Regulates Transcriptional Interference and Gene Repression": Table S6

**Supplemental Table 7. Plasmids generated in this study**

| **Plasmid Number** | **Plasmid Info** |
| --- | --- |
| pUB1274 | *HIS3-plexO:HIS3LUTI* |
| pUB1645 | *URA3-pTEF1:ADE2LUTI* |
| pUB1577 | *LEU2-pSWI3:SWI3* |
| pUB1578 | *LEU2-pSWI3:swi3-E815X* |
| pUB2025 | *LEU2-pSNF2:SNF2* |
| pUB2050 | *LEU2-pSNF2:snf2-W935R* |
| pUB2049 | *LEU2-pSNF2:snf2-Q928K* |
| pUB2204 | *TRP1-HNT1-3V5* |
| pUB2151 | *TRP1-HNT1(LUTIΔ)-3V5* |
| pUB2462 | *TRP1-HNT1(LUTI-CYC1t)-3V5* |
| pUB2479 | *TRP1-ADI1-3V5* |
| pUB2480 | *TRP1-ADI1(pDISTΔ)-3V5* |
| pUB2481 | *TRP1-ODC2-3V5* |
| pUB2482 | *TRP1-ODC2(pDISTΔ)-3V5* |
| pUB2485 | *TRP1-ERG27-3V5* |
| pUB2486 | *TRP1-ERG27(pDISTΔ)-3V5* |
